# Role of Dynamin-Related Proteins 2 and SH3P2 in Clathrin-Mediated Endocytosis in Plants

**DOI:** 10.1101/2023.10.09.561523

**Authors:** Nataliia Gnyliukh, Alexander Johnson, Marie-Kristin Nagel, Aline Monzer, Annamaria Hlavata, Erika Isono, Martin Loose, Jiří Friml

## Abstract

Clathrin-mediated endocytosis (CME) is vital for the regulation of plant growth and development by controlling plasma membrane protein composition and cargo uptake. CME relies on the precise recruitment of regulators for vesicle maturation and release. Homologues of components of mammalian vesicle scission are strong candidates to be part of the scissin machinery in plants, but the precise roles of these proteins in this process is not fully understood. Here, we characterised the roles of Plant Dynamin-Related Proteins 2 (DRP2s) and SH3-domain containing protein 2 (SH3P2), the plant homologue to Dynamins’ recruiters, like Endophilin and Amphiphysin, in the CME by combining high-resolution imaging of endocytic events *in vivo* and characterisation of the purified proteins *in vitro*. Although DRP2s and SH3P2 arrive similarly late during CME and physically interact, genetic analysis of the D*sh3p1,2,3* triple-mutant and complementation assays with non-SH3P2-interacting DRP2 variants suggests that SH3P2 does not directly recruit DRP2s to the site of endocytosis. These observations imply that despite the presence of many well-conserved endocytic components, plants have acquired a distinct mechanism for CME.

**One Sentence Summary:** In contrast to predictions based on mammalian systems, plant Dynamin-related proteins 2 are recruited to the site of Clathrin-mediated endocytosis independently of BAR-SH3 proteins.

## Introduction

Clathrin-mediated endocytosis (CME) is a crucial cellular process that enables cell response to changes in the extracellular environment by internalizing plasma membrane (PM), small molecules, and transmembrane proteins. During CME, cargo is encapsulated within clathrin-coated vesicles (CCVs), which subsequently detach from the PM to undergo further trafficking, and subsequent cargo recycling or degradation (McMahon and Boucrot, 2011). CME plays a vital role in various cellular functions in plants, including cell wall synthesis, nutrient uptake, immune response and hormone signalling (Barberon et al., 2011; Bashline et al., 2013; Chen et al., 2011; Dhonukshe et al., 2007; Irani et al., 2012; Luschnig and Vert, 2014; Mbengue et al., 2016; Narasimhan et al., 2021; Paciorek et al., 2005; Pan et al., 2009; Postma et al., 2016; Sánchez-Rodríguez et al., 2018; Wang et al., 2017). CME regulates the internalization of important PM proteins such as PIN-FORMED (PIN), Brassinosteroid Insensitive 1 (BRI1), borate receptor (BOR1), iron-regulated transporter 1 (IRT1), and cellulose synthase A (CESA) (Barberon et al., 2011; Bashline et al., 2013; Claus et al., 2018; Dhonukshe et al., 2007; Di Rubbo et al., 2013; Yoshinari et al., 2016; Zhang et al., 2019). Our understanding of CME mechanism in plants originates mainly from mechanistic predictions in mammalian and yeast cells, as plants (mainly the model *Arabidopsis thaliana*) possess homologous proteins to most key CME components including clathrin heavy chain (CHC) and clathrin light chain (CLC), adapter proteins (AP), and Dynamin-Related Proteins (DRPs) (Dhonukshe et al., 2007; Gadeyne et al., 2014; Lu et al., 2016; McMahon and Boucrot, 2011). This led to the belief that plant CME works analogous to other eukaryotic systems, although the physiological and mechanistic properties of plant cells are strikingly different from those of mammals and yeasts (Heidstra and Sabatini, 2014). The presence of a rigid cell wall and central vacuole creates high turgor pressure that influences mechanisms of various cell processes, including CME (Chen et al., 2011). Still, unlike in yeasts, in plants actin is not present at the site of endocytic vesicle formation (Narasimhan et al., 2020). Instead, a TPLATE complex has been suggested to be a plant-specific driver of membrane invagination (Johnson et al., 2021). These and other studies highlight significant differences between CME in plants and mammals and question its conserved mechanism (Backues et al., 2010; Fujimoto et al., 2010; Gadeyne et al., 2014). Therefore, to understand the mechanism of plant endocytosis, further investigation is needed.

During the final stage of CME, after CCV has been fully formed, scission machinery mediates the release of the vesicle from PM (McMahon and Boucrot, 2011). In mammalian systems, essential role in this process is played by a large GTPase dynamin (Marks et al., 2001). It consists of GTP hydrolysis (GTPase) domain, Middle domain, GTPase effector domain (GED), and membrane binding Pleckstrin homology (PH) domains followed by protein-protein interaction Proline-rich domain (PRD). This large GTPase assembles into oligomers around a highly curved membrane connecting vesicle to PM and releases the vesicle via GTP-hydrolysis-based conformational changes of the oligomer (Antonny et al., 2016). Thus, dynamin plays an important role in synaptic vesicle recycling, receptor-mediated endocytosis sequestering ligands into invaginated coated pits (Perrais, 2022; Prichard et al., 2022). In yeasts, Vps1, Dnm1 and Mgm1 have been described as homologues of dynamin, which possess similar properties and function in CME (Lee et al., 2017; Rooij et al., 2010). They consist of the N-terminal GTPase domain, Middle domain and GED, but lack a PH domain and the canonical PRD. However, unlike mammalian dynamins, yeast homologues have been shown to play a role not only in vesicle fission, but also for the invagination and correct morphology of cortical actin patches (Rooij et al., 2010b; Yu and Cai, 2004). In plants, many potential DRPs have been identified based on sequence homology and found to play a role in mitochondrial and chloroplast fission, during cytokinesis, cell plate formation and CME (Fujimoto et al., 2010, 2010; Lam et al., 2002). Specifically, members of the DRP1 and DRP2 subfamilies were shown to co-localize with other endocytic markers on the PM. While members of the DRP1 family have a domain organization similar to their yeast homologues (lacking PH domain and PRD), only DRP2s have the same domain organization as their mammalian counterpart (Hong et al., 2003). They co-localize with CLC and proteins of the DRP1 subfamily on the PM (Fujimoto et al., 2010). Two members of the DRP2 subfamily, DRP2A and DRP2B, show a great sequence similarity and have previously been shown to be functionally redundant (Backues et al., 2010). Based on these observations, DRP2s have been thought to be involved in vesicle scission but has not de definitely examined.

In mammals and yeast, the recruitment of dynamin to the high-curved membrane neck connecting the vesicle and the PM was found to be facilitated by bin/amphiphysin/Rvs (BAR) domain-containing proteins like endophilin (Endo2) and amphiphysin (Amph1) (Bhatia et al., 2009; Gallop et al., 2006; Pant et al., 2009; Renard et al., 2015). The BAR domain was shown to recognize high membrane curvature, while the src homology-3 (SH3) domain interacts with other signalling and regulatory proteins (Peter et al., 2004; Xin et al., 2013). During membrane scission, the SH3 domain of Endo2 and Amph1 interacts with the PRD of dynamin, recruiting it to the site of the CME (Luo et al., 2016; Sundborger et al., 2014). Additionally, BAR domain-containing proteins can deform membranes, potentially aiding scission (Farsad et al., 2001; Peter et al., 2004). Disruption of these proteins’ function severely impairs synaptic vesicle endocytosis at central nerve terminals (Jockusch et al., 2005; Kontaxi and Cousin, 2023; Shupliakov et al., 1997). Altogether, studies in mammalian cells revealed the identity and regulation of several proteins required for vesicle formation during CME. In yeasts, although Vps1 lacks the canonical PRD, its interaction with the Amph1-homologue Rvs167 is important for vesicle scission, suggesting some degree of similarity between the mammalian and yeast systems (Smaczynska-de Rooij et al., 2012). Importantly, these studies emphasized the importance of BAR-domain-containing proteins for this process. However, if this mechanism is conserved in plants is yet to be clarified.

Three members of plant BAR-SH3-domain-containing proteins (SH3P1, SH3P2 and SH3P3) in Arabidopsis were previously studied during cell plate assembly, endosomal sorting, intracellular trafficking, and autophagosome biogenesis (Ahn et al., 2017; Baquero Forero and Cvrčková, 2019; Kolb et al., 2015; Nagel et al., 2017; Zhuang et al., 2013; Zhuang and Jiang, 2014). The importance of SH3P2 for vesicle trafficking (interaction with ESCRT-I/III system and ubiquitinated proteins in CCVs), autophagosome formation, and cell plate formation has been demonstrated; however, little was done to understand its role in the process of vesicle release from PM. Some studies have suggested involvement of SH3Ps in plant CME, where it hypothesised that members of the SH3P family recruit the scission machinery to the clathrin-coated pit (CCP), similar to their mammalian homologues such as Endo2 and Amph1 (Lam et al., 2001; Lebecq et al., 2022). For instance, SH3P proteins have been found to localize on clathrin-positive vesicles and co-localizes with Auxilin-LIKE1, which likely participates in the uncoating of CCVs (Adamowski et al., 2022; Dahhan et al., 2022; Nagel et al., 2017). Notably, the SH3 domain of SH3P3 interacts with members of the DRP2 family (Lam et al., 2002). Additionally, the BAR domain of SH3P2 has been shown to bind and tubulate liposomes *in vitro* (Ahn et al., 2017). While these findings suggest involvement of SH3P proteins in plant CME, a comprehensive characterization of their behaviour and properties *in vivo* and *in vitro* is currently missing. This lack of knowledge makes it difficult to propose a mechanism for plant CME and the roles of the proteins involved.

In this study, we aimed to obtain new insights into plant CME, focusing specifically on the roles of DRP2s and SH3P2. By using high-resolution imaging *in vivo,* we found that SH3P2, similar to DRP2A and DRP2B, arrives at the end of the endocytic event. In *in vitro* experiments, we found that purified SH3P2 is able to deform membranes as previously described for mammalian homologues. Our data further reveal co-localization *in planta* and direct interaction between purified SH3P2 and DRP2B *in vitro*. We found that PM internalization is significantly impaired in the Δ*sh3p1,2,3* triple mutant, while DRP2A, CLC2, and TPL endocytosis markers show normal dynamics. Thus, our findings shed light on the dynamics of two key players in plant endocytic scission machinery, suggesting that DRP2s have a distinct recruiting mechanism from its mammalian counterparts, largely independent of SH3P2.

## Results

### DRP2 proteins arrive at the end of the endocytic event

The role of different proteins involved in endocytosis show a stereotypical timing of recruitment depending on their function (McMahon and Boucrot, 2011). For example, regulators involved in vesicle scission, like dynamin, Endo2, and Amph1, are recruited right before the vesicle is pinched off and released from the PM (Rosendale et al., 2019; Taylor et al., 2011). Given the homology of DRP2B to dynamins, and that it previously found to co-localise with CLC at the PM (Fujimoto et al., 2010), we decided to obtain more information on their dynamics together with CME markers *in planta* at high spatiotemporal resolution to gain insights into role of DRP2s in plant endocytosis.

We used Total internal reflection fluorescence microscopy (TIRF-M) of epidermal root cells, which allowed us to create high-resolution time-lapse images of protein dynamics exclusively on the PM (Johnson et al., 2020; Trache and Meininger, 2008). Before imaging, we confirmed that DRP2A tagged with C-terminal GFP was functional by complementing gametophyte lethal phenotype of *drp2a-/-;drp2b-/-* double mutant (Fig. S1) (Backues et al., 2010). TIRF-M of DRP2A-GFP in *drp2a-/-;drp2b-/-* showed increase in dynamics and spot density compared to the control (DRP2A-GFP in *drp2a-/-)*, potentially indicating a compensatory mechanism for lack of DRP2B (Fig. S1E-G).

To analyse the recruitment of DRP2A exclusively within CME events, we simultaneously visualised DRP2A-GFP and CLC2 fluorescently tagged with tagRFP, a CME marker, and applied an automated analysis to obtain the precise arrival and departure times of the protein (Fig. 1A-B) (Johnson et al., 2020). Our data showed that 60% of CLC2 foci co-localised with DRP2A (Fig. 1F-G), suggesting that a significant amount of CME foci include DRP2A. Analysis of fluorescent profiles of DRP2A co-localised with CLC2-tagRFP showed that the maximum intensity of DRP2A recruitment occurred before the CLC2 signal disappeared, marking the departure of the vesicle from PM (Fig. 1C). Average lifetime of these events was 46.3±0.22 s with a density of 39.64±1.68 spots per region of interest [640 µm^2^] (ROI)^−1^, compared to a long population of endocytosis events represented by TPLATE (TPL) and CLC2 (∼43 s) (Fig. 1D-E) (Narasimhan et al., 2020). We observed similar results for the second member of the DRP2 family, DRP2B-GFP with CLC2-tagRFP (Fig. S2).

**Fig. 1.**
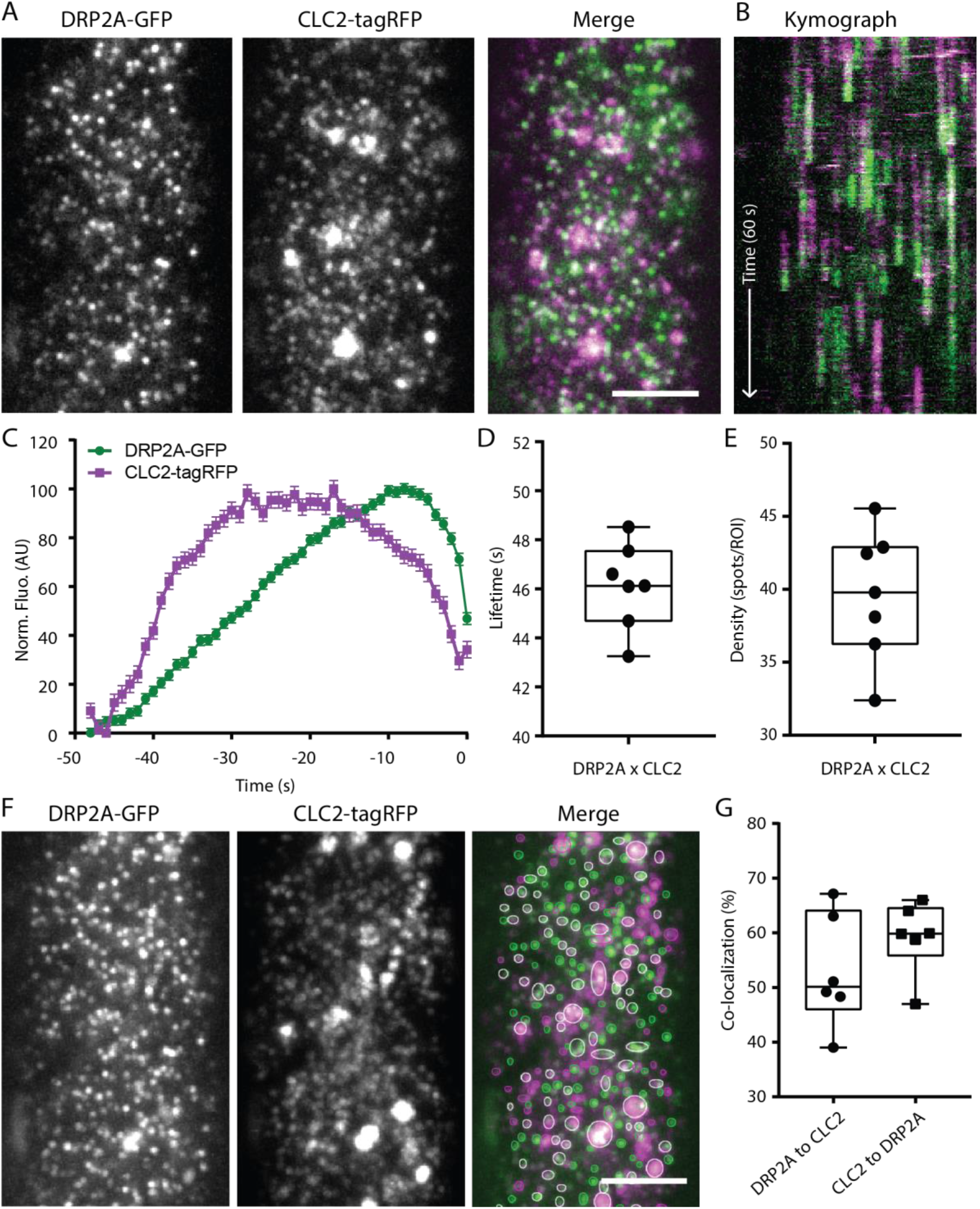
Recruitment of DRP2A to the site of vesicle formation. (A) TIRF-M images of a cell surface of root epidermal cell expressing fluorescently tagged DRP2A-GFP and CLC2-tagRFP. (B) Representative kymograph of DRP2A and CLC2 lifetimes on the PM. Scale bar: 5 µm. (C-E) Data from seven independent experiments were combined to generate a (C) mean recruitment profile of DRP2A to the site of endocytosis, (D) mean lifetime, 46.12±0.6 s, and (E) mean density 39.63±1.6 spots ROI^−1^ of CME events. Plots indicate Mean±SEM, n=7 cells from independent roots, 22,432 tracks. (F) Representative image of co-localisation analysis of DRP2A and CLC2 foci. Scale bar: 5 µm. (G) Quantification of co-localised spots. 52.98±4.2 % of DRP2A were co-localised to CLC2, and 59.25±2.7% of CLC2 were co-localised to DRP2A.

The observed recruitment profile of DRP2A is typical for proteins involved in the late stages of CCV formation, such as during scission. Therefore, these observations support the hypothesis that DRP2 proteins are part of the CME scission machinery.

### SH3P2 arrives at the end of endocytosis, similar to DRP2

Recruitment of dynamin and Vps1 are tightly linked to the arrival of Amph1/Endo2 and Rvs167 to the site of CCV formation in mammals and yeast, respectively. Previously, plant homologue of these proteins SH3P2 was shown to co-localize with CLC and co-fractionation with CCVs, providing initial indications for its function in CME (Adamowski et al., 2022; Dahhan et al., 2022; Nagel et al., 2017).

To investigate the function of SH3P2 in plant CME, we quantified the recruitment of SH3P2-sGFP compared to CLC2-mOrange used as a reference for CME events. The co-localization frequency of CLC2 and SH3P2 at a given time point was 60% of the CLC2 similar to what we had observed for DRP2A co-localization with CLC2 (Fig. 2F-G). Next, we analysed individual endocytosis events positive in both CLC2 and SH3P2. The peak of the SH3P2-sGFP signal occurred just before the drop of the CLC2-mOrange signal, indicating of CCV scission (Fig. 2A-C). The average lifetime of SH3P2- and CLC2-positive events was 40.54±0.19 s with an average density of 44.26±3.4 spots ROI^−1^ (Fig. 2C-E), which is in line with data obtained for DRP2s and lifetime of long population of endocytosis events reported in Narasimhan et al., 2020. The profile of SH3P2 recruitment to the site of CCV formation suggests that SH3P2 functions at the final stage of CME in plants.

**Fig. 2.**
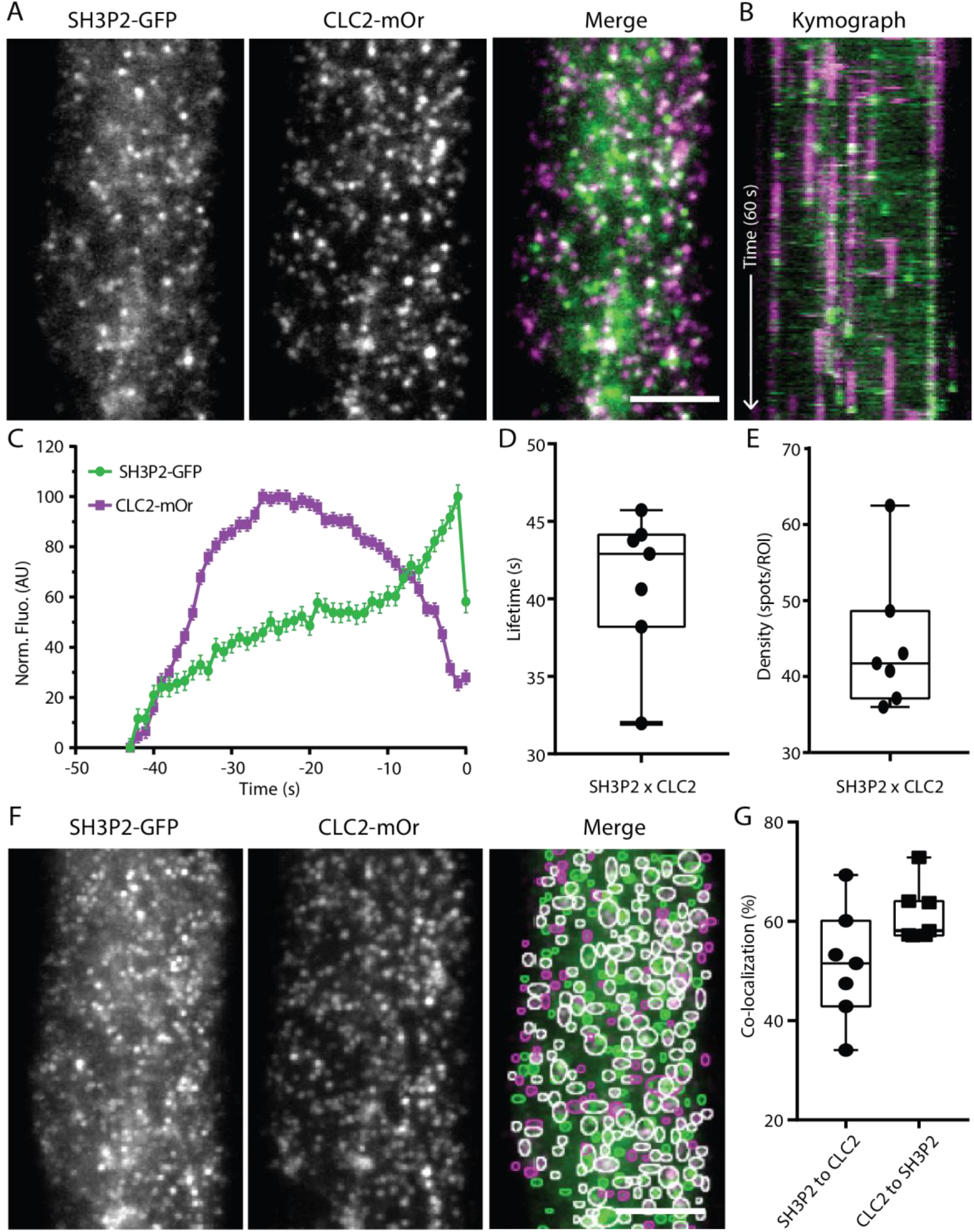
SH3P2 arrives at the end of the endocytosis event. A) TIRF-M images of a cell surface of root epidermal cell expressing fluorescently tagged SH3P2-sGFP and CLC2-mOrange. Scale bar: 5 µm. (B) Representative kymograph of SH3P2 and CLC2 lifetimes on the PM. (C-E) Data from seven independent experiments were combined to generate a (C) mean recruitment profile of SH3P2 to the site of endocytosis, (D) mean lifetime 41.05±0.5 s, and (E) mean density 44.26±3.4 spots ROI^−1^ of CME events. Plots indicate Mean±SEM, n=7 cells from independent roots, 24,170 tracks. (F) Representative image of co-localisation analysis of SH3P2 and CLC2 foci. Scale bar: 5 µm. (G) Quantification of co-localised spots. 51.25±4.33 % of SH3P2 were co-localised to CLC2, and 61.49±2.22 % of CLC2 were co-localised to SH3P2.

These *in vivo* observations revealed that both SH3P2 and DRP2A are specifically recruited at the end of the CME events on the PM, much like their homologues during CME in other systems.

### SH3P2 binds and bends membranes *in vitro*

Complementation of protein dynamics in live cells with *in vitro* experiments of individual proteins provides a more detailed and controlled understanding of their function. Therefore, to study the properties of DRP2s and SH3P2, we decided to test binding, bending and fission abilities using purified proteins *in vitro*. Unfortunately, after extensive efforts, we were not able to purify GTPase active full-length DRP2A or DRP2B. It is known that GTPase activity is crucial for its function *in vivo*, and therefore we continued the characterisation of SH3P2 protein alone.

In mammalian systems, the BAR-SH3-domain-containing proteins Endo2 and Amph1, bind to and remodel membranes, which is crucial for the successful vesicle formation (Blood and Voth, 2006; Habermann, 2004). Previous studies have demonstrated that the isolated BAR domain of *At*SH3P2 predominantly binds negatively charged membranes and induces vesicle deformation *in vitro* (Ahn et al., 2017).

To assess the membrane binding and remodelling capacity of SH3P2, we purified the bacterially expressed protein (Fig. S3A). Mass photometry measurements showed two peaks in the histogram corresponding to the mass of the SH3P2 monomer (∼39 kDa) and dimer (∼78 kDa) (Young et al., 2018). With an increase of protein concentration in solution from 75 nM to 100 nM, we observed an increase percentage of dimers (60% vs 85%), suggesting that dimerization is concentration dependent (Fig. S3B). This confirms that purified SH3P2, similar to BAR-domain-containing proteins from mammalian system, can dimerize (Jhaveri et al., 2021; Youn et al., 2010).

Next, we wanted to understand the ability of SH3P2 to bind membranes, as in mammalian systems, the cooperative membrane binding of BAR-domain containing proteins and dynamin is important for vesicle release from the PM (Meinecke et al., 2013). Therefore, we conducted a liposome sedimentation assay, where the protein of interest only sediments when bound to phospholipid vesicles. As it has been shown that PM sites of occurring CME are enriched with negatively charged lipids (Martin, 2001), we decided to study the lipid binding preferences of SH3P2. We generated large unilamellar vesicles (LUVs) of different lipid compositions, with and without negatively charged lipids, such as phosphatidic acid (PA) and phosphatidylinositol 4,5-bisphosphate (PI(4,5)P_2_). We found that SH3P2 exhibited a preference for binding to LUVs containing PA and even more PI(4,5)P_2_ compared to LUVs with only 1,2-Dioleoyl-sn-glycero-3-phosphocholine (DOPC) or DOPC mixed with 1,2-dioleoyl-sn-glycero-3-phospho-L-serine (DOPS) (Fig. S3C-D). Finally, to test the membrane deforming capability of SH3P2, we incubated the purified proteins with LUVs with DOPC/DOPS/PI(4,5)P_2_ and performed Transmission Electron Microscopy (TEM) experiments. Compared to the control vesicles without protein, LUVs showed a significant percentage of membrane deformation in presence of SH3P2 (Fig. 3), demonstrating the ability of the protein to deform membranes.

**Fig. 3.**
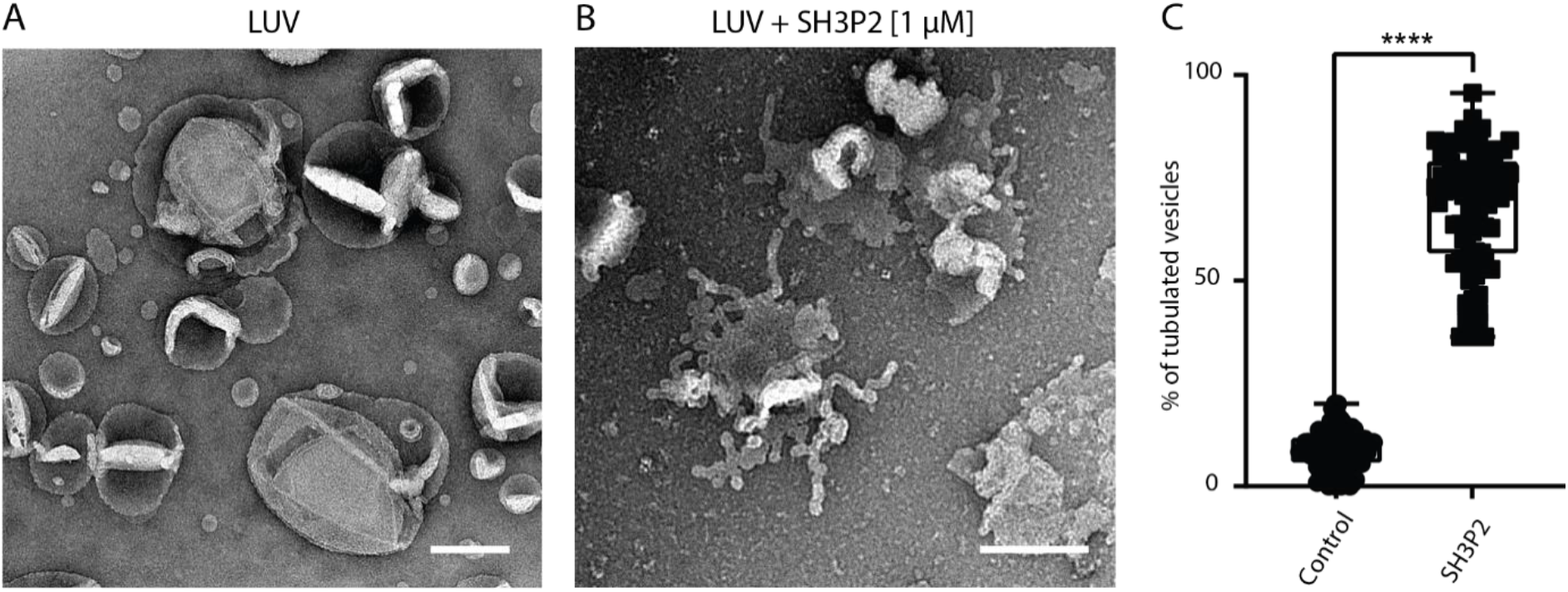
Full-length SH3P2 protein bends membranes *in vitro*. (A) Example TEM overviews of LUVs after 5 min incubation in control conditions or (B) with 1 μM SH3P2. Scale bar, 200 nm. (B) Quantification of the percentage of LUVs displayed tubulation. 8.83% of LUVs displayed tubulation in control conditions. 68.02% of LUVs incubated with SH3P2 displayed tubulation. N, control, 77 images; SH3P2, 85 images pooled from three independent experiments. Plot, Mean ± SEM, ****P < 0.001, t-test to compare to control.

In conclusion, reconstitution experiments *in vitro* showed that FL SH3P2, similarly to its mammalian homologues, dimerizes and preferentially binds negatively charged lipids. Moreover, similarly to BAR domain of SH3P2 alone, full-length SH3P2 promotes tubulation of LUVs *in vitro*.

### DRP2A and SH3P2 interact *in vitro* and co-localize *in vivo*

Since recruitment profiles of SH3P2 and DRP2s indicate their involvement in the terminal stages of the CME event, we further investigated the spatiotemporal relation between these proteins. Taking into account that their recruitment is similar to that of mammalian dynamin and BAR-domain proteins (Taylor et al., 2011), we hypothesised alike interaction mechanism between SH3P2 and DRP2s. While mammalian dynamin interacts with other proteins through its PRD of 13 proline-rich motifs (PRMs) (Okamoto et al., 1997), DRP2s contain only two highly conserved PRMs localised at the beginning of GED domain and in the middle of PRD, respectively (DRP2A:PRM1 – RKPIDPEE and PRM2 – RLPPAPPPTG; DRP2B: PRM1 – RKPVDPEE, PRM2 - RLPPAPPQS) (Fig. 4A) (Hong et al., 2003; Schmid and Frolov, 2011).

**Fig. 4.**
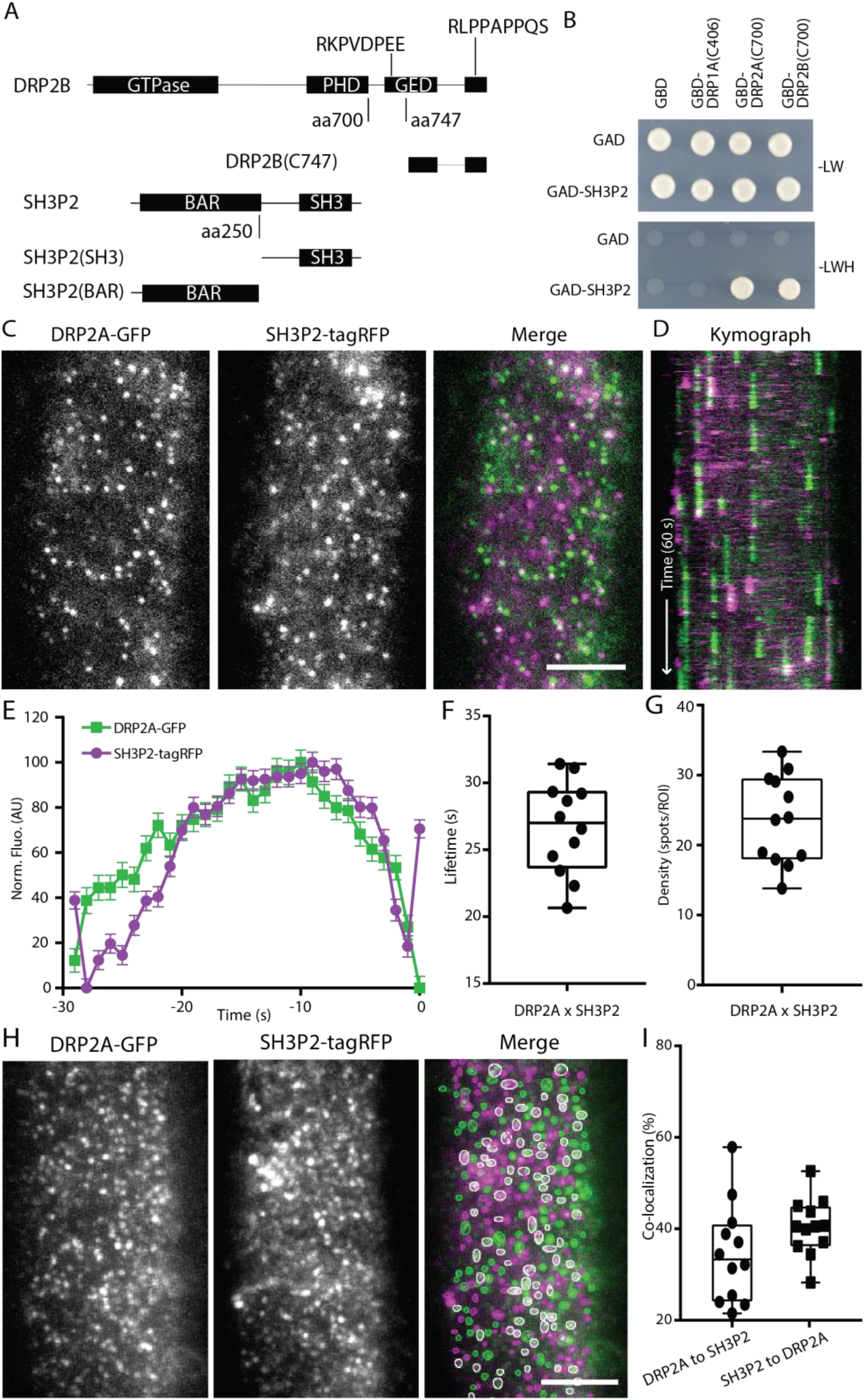
DRP2A and SH3P2 interact and co-localize. (A) Schematic presentation of full-length DRP2B and SH3P2 the truncated constructs DRP2B(C747), SH3P2(SH3) and SH3P2(BAR). (B) YTH analyses of GAD-SH3P2 with GBD-fusions of DRP2A and DRP2B. Yeast transformants were grown on media lacking leucine and tryptophan (−LW) or leucine, tryptophan, and histidine (−LWH) supplemented with 1 mM 3-amino-1,2,4-triazole (3-AT) to test their auxotrophic growth. Empty vector GBD and a truncated version of DRP1A were used as negative controls. The expression of all fusion proteins was verified by immunoblotting shown in Fig. S4. (C) TIRF-M images of a cell surface of root epidermal cell expressing fluorescently tagged DRP2A-EGFP and SH3P2-tagRFP. Scale bar: 5 µm. (D) Representative kymograph of DRP2A and SH3P2 lifetimes on the PM. (E-G) Data from twelve independent experiments were combined to generate an (E) mean recruitment profile of SH3P2-positive DRP2A foci, (F) mean lifetime 26.68±0.5 s, and (G) mean density 23.63±1.8 spots ROI^−1^ of CME events. Plots indicate Mean±SEM, n=12 cells from independent roots, 16,916 tracks. (H) Representative image of co-localisation analysis of SH3P2 and DRP2A foci. Scale bar: 5 µm. (I) Quantification of co-localised spots. 34.58±3.1% of DRP2A were co-localised to SH3P2, and 40.41±1.7% of SH3P2 were co-localised to DRP2A.

First, we tested the interaction between SH3P2 and two C-terminal parts of DRP2 proteins, both containing PRMs, using Yeast-Two-Hybrid (YTH) assay. As a negative control, we used C-terminal part of DRP1A protein, a member of DRP subfamily 1 that lacks both PH domain and PRD. Our results showed that C-terminal parts of DRP2A and DRP2B, that contain both PRMs, were able to interact with SH3P2 in YTH (Fig. 4B, Fig. S4A). Expectedly, we could not detect any interaction between SH3P2 and DRP1A. Together, these data suggest that SH3P2 interacts with the C-terminus of DRP2 and this interaction could be mediated through the PRMs in DRP2A and DRP2B.

To further investigate the relevance of our *in vitro* observations for CME, we generated plant line containing SH3P2-tagRFP and DRP2A-GFP markers to study their dynamics and co-localization *in vivo* (Fig. 4C-D). We were able to detect foci that were spatiotemporally positive in both channels and the average lifetime of these events was 26.6±0.14 s (Fig. 4E-G). This lifetime is almost twice shorter compared to the lifetime of DRP2A and SH3P2 with CME marker (∼46 s and ∼41 s, respectively) (Fig. 1D, 2D), which reflects their arrival during the late stage of CME. The fluorescent profiles of DRP2A and SH3P2 suggest simultaneous peak of their arrival, although DRP2A seems to arrive slightly earlier than SH3P2 (Fig. 4E). The average density of these events was 23.63±1.82 spots ROI^−1^, which was lower than foci density of DRP2A and SH3P2 with CLC2 (∼40 and ∼44 spots ROI^−1^) (Fig. 4G). We further checked the percentage of co-localized SH3P2-tagRFP and DRP2A-GFP foci (Fig. 4H) and found that a frequency of 34.6% for SH3P2-tagRFP and DRP2A-GFP, whereas 40.41% of DRP2A-GFP was co-localized to SH3P2-tagRFP at a given time point (Fig. 4I). These data suggests the existence of CME events that have either DRP2A or SH3P2 separately. However, the events that have both SH3P2 and DRP2A arriving at the PM together may potentially function together, similarly to mammalian Endo2/Amph1 and dynamin.

Next, we determined the importance of the DRP2 PRMs for the interaction with the SH3 domain of SH3P2 (Fig. 5A). Minimal domain analyses using YTH showed that PRM1 cannot interact with SH3 domain of SH3P2 alone, whereas PRM2 showed interaction with both SH3P2 full-length protein and C-terminal fragment containing the SH3 domain (Fig. 5B). The N-terminus of SH3P2 containing the BAR domain did not interact with either PRMs. Moreover, our results show that PRM1 cannot interact with SH3 domain of SH3P2 alone (Fig. 5B). In contrast, the presence of the PRM2 motif, located in PRD, alone was sufficient for the protein-protein interaction (Fig. 5B, Fig. S4B-C).

**Fig. 5.**
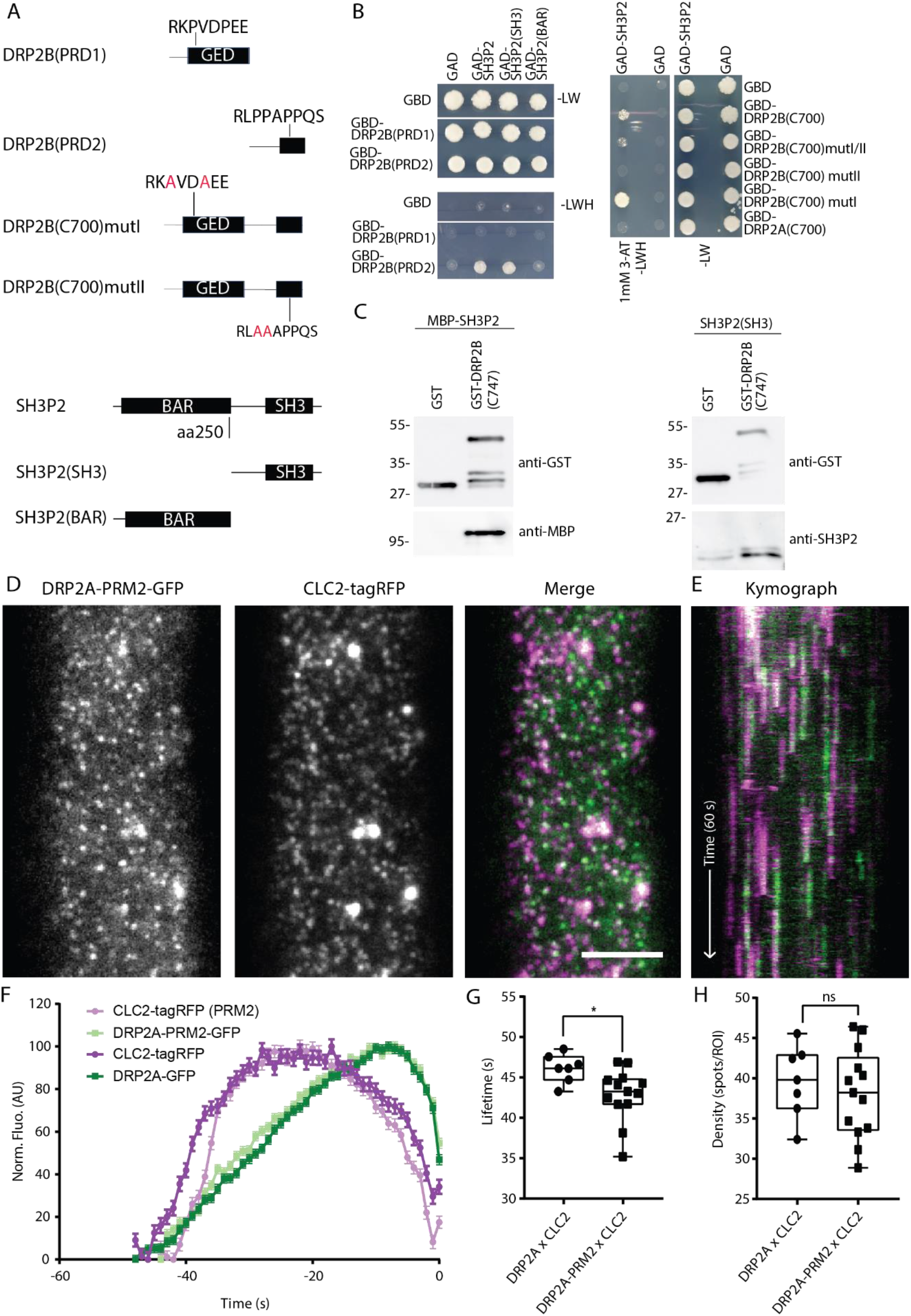
PRM2 is important for interaction with SH3P2 *in vitro*, but not *in vivo*. (A) Schematic presentation of the truncated constructs DRP2B (C747), DRP2B (PRD1) and DRP2B (PRD2), DRP2B (C700)mutI and DRP2B(C700)mutII, SH3P2 (SH3) and SH3P2 (BAR). (B) YTH analyses of GAD-SH3P2 with GBD-fusions of truncated versions of DRP2B. Yeast transformants were grown on media lacking leucine and tryptophan (−LW) or leucine, tryptophan, and histidine (−LWH) supplemented with 1 mM 3-amino-1,2,4-triazole (3-AT) to test their auxotrophic growth. Empty vectors, GAD and GBD, were used as negative controls. The expression of all fusion proteins was verified by immunoblotting shown in Fig. S4. (C) In vitro-binding assay of SH3P2 with DRP2B(C747). GST was used as a negative control. Bead-bound GST and GST-DRP2B(C747) were incubated with equal amounts of full-length MBP-SH3P2. After intensive washing, bead-bound materials were subjected to immunoblotting using anti-MBP and anti-GST antibodies. (D) TIRF-M images of a cell surface of root epidermal cell expressing fluorescently tagged DRP2A-PRM2-EGFP and CLC2-tagRFP. Scale bar: 5 µm. (E) Representative kymograph of DRP2A-PRM2 and CLC2 lifetimes on the PM. (F-H) Data from nine independent experiments for DRP2A-PRM2 x CLC2 were combined and compared to DRP2A x CLC2 from Fig. 1, to generate an (F) mean recruitment profile of DRP2A and DRP2A-PRM2 foci, (G) mean lifetime 43.72±0.18 s, and (H) mean density 40.05±1.76 spots ROI^−1^ of CME events. Plots indicate Mean±SEM, DRP2A-PRM2 x CLC2 n=9 cells from independent roots, 29,407 tracks. Plot, Mean ± SEM, ns >0.05, t-test to compare to control.

We further designed point mutations in one or both PRMs and tested their interaction with SH3P2 using YTH assays (Fig. 5A). To reduce autoactivation of the GAL4-binding domain, the selection medium was supplemented with 3-amino-1, 2, 4-triazole (3-AT). When both RPMs were mutated, we did not detect any interaction between DRP2B and SH3P2. The presence of the PRM2 motif alone was sufficient for the protein-protein interaction. Mutating PRM2 prevented the interaction with SH3P2, whereas mutation in PRM1 did not (Fig. 5B). The direct interaction between SH3P2 and PRM2 of DRP2B was confirmed by *in vitro* pull-down assay, using GST-DRP2B(C747), full-length protein MBP-SH3P2 and untagged SH3P2(SH3) (Fig. 5C). Summarising these data, we confirmed that SH3P2 can interact with both members of the DRP2 subfamily and that only PRM2 of DRP2B is crucial for the interaction with SH3 domain of SH3P2.

To complement these *in vitro* interaction experiments, we generated plant CLC2-tagRFP lines containing DRP2A-GFP with the point mutation in the PRM2 (RL**AA**APP**QS**). We analysed the dynamics of DRP2A-PRM2-GFP × CLC2-tagRFP compared to WT DRP2A-EGFP × CLC2-tagRFP using TIRF-M (Fig. 5D-E). Despite the presence of point mutation in this interaction site, we were able to detect around 60% of CLC2 foci co-localized with DRP2A-PRM2 with dynamics of both proteins similar to the WT DRP2A (Fig. 5F, S5). The mean lifetime of DRP2A-PRM2-EGFP × CLC2-tagRFP was reduced by 3 s (43.72±0.1 s) compared to the control (46.12±0.6 s), but the foci density remained the same (Fig. 5G-H). Additionally, we generated a similar mutation in PRM2 of DRP2B (RL**AA**APPQS). No significant change was detected in either the lifetime or foci density of DRP2B-PRM2 mutant compared to WT DRP2B, showing that DRP2B was not affected by the PRM2 mutation (Fig. S6). Overall, although we detected a reduced lifetime of DRP2A-PRM2-CLC2 foci, these proteins had a dynamic behaviour at the PM, indicating that despite interaction mutation DRP2s were efficiently recruited to the site of CCV formation.

Additionally, we tested CLC2 dynamics in case of overexpression of SH3P2 SH3 domain without membrane binding domain. The abundance of SH3 domain in the cytosol should sequester interacting proteins, like DRP2s, to prohibit them from recruitment to the PM, causing impairment of CME (Gad et al., 2000; Szaszák et al., 2002). To test this, we generated a line with a CLC2-GFP marker and overexpressed the SH3P2 SH3 domain tagged with mCherry marker or free mCherry as a control (Fid. S7A-D). Analysis of CLC2 foci dynamics on PM showed no difference in neither lifetime nor density (Fig. S7E-G).

To conclude, we identified the PRM of DRP2s to be involved in protein-protein interaction with SH3P2 *in vitro*; however, the mutation in this motif did not significantly affect endocytosis or DRP2 dynamics *in vivo* revealing no functional consequence of the *in vitro* interaction.

### *Δsh3p1,2,3* shows normal DRP2A, CLC2 or TPLATE dynamics

In order to further investigate role of SH3P proteins within plant CME, we used a new mutant of all three proteins of the SH3P family. It was generated by combining CRISPR-Cas technique for SH3P1 and SH3P2 and T-DNA insertion line of SH3P3 (Adamowski et al., 2022). The D*sh3p1,2,3* triple mutant was reported to have defects in plant growth and development, as well as seed germination. These phenotypes are more pronounced than one reported for various combinations of T-DNA alleles (Ahn et al., 2017; Nagel et al., 2017). On the contrary, the SH3P2 RNA interference (RNAi) line displayed severe defects in seedling development, showing the importance of SH3P2 alone for plant development (Zhuang et al., 2013).

We used D*sh3p1,2,3* triple mutant line to investigate the potential impairment of CME and defects in CME marker and cargo dynamics. Prior to the visualisation of CME marker in this line, we assessed the general membrane uptake of PM in root epidermal cells by measuring the uptake of the non-permeable membrane dye FM4-46 (Bolte et al., 2004; Jelínková et al., 2019). This assay allows testing membrane internalisation by analysis of formation of intracellular vesicles, stained with FM4-64. We observed a significant reduction of membrane uptake in root cells of D*sh3p1,2,3* triple mutant compared to the WT, D*sh3p1,2* double and D*sh3p3* single mutants, which all exhibited a normal rate of membrane internalisation (Fig. S8A-B). This is a significant impairment of PM internalisation; however, this does not directly represent the rate of CME in plant cells. Therefore, we further tested if mutation of SH3P proteins influences the recruitment of DRP2A to the site of CCV formation.

To do so we studied the dynamics of DRP2A-GFP in D*sh3p1,2,3* triple mutant background compared to the control DRP2A-GFP (Fig. 6A-D). Quantitative analysis of DRP2A lifetime persistence revealed that there was no significant difference between control and mutant plants (Fig. 6E-F). Also, no change in density of DRP2A foci was detected (Fig. 6G). These data contradict the hypothesis that SH3P proteins are crucial for the recruitment of DRP2s to the site of endocytosis, as DRP2A was still dynamically appearing and disappearing on the PM. To further investigate if D*sh3p1,2,3* triple mutation influences the dynamics of other CME markers we analysed the lifetime of total CLC2 events on PM. Previously, it was shown that the CLC2 density in the D*sh3p1,2,3* background was reduced compared to control, although no measurement of lifetime or recruitment profiles was done (Adamowski et al., 2022). Therefore, using the unbiased high throughput analysis we provide a robust measurement of CLC2-GFP dynamics in D*sh3p1,2,3* background and saw a 1,7 s increase in lifetime in CLC2 D*sh3p1,2,3*; however, no difference in density in the mutant (Fig. S9). To then analyse *bona fide* CME events, and exclude unsuccessful and aborted events, in the D*sh3p1,2,3* triple mutation we visualize the dynamics of TPL × CLC2 positive foci and compared it to the control line (Fig. 6H-I). Quantitative analysis of the recruitment profiles of TPL and CLC2 positive events revealed no significant difference between control and mutant (Fig. 6J-K). Density of TPL × CLC2 was only slightly increased in D*sh3p1,2,3* mutant background compared to the control (Fig. 6L).

**Fig. 6.**
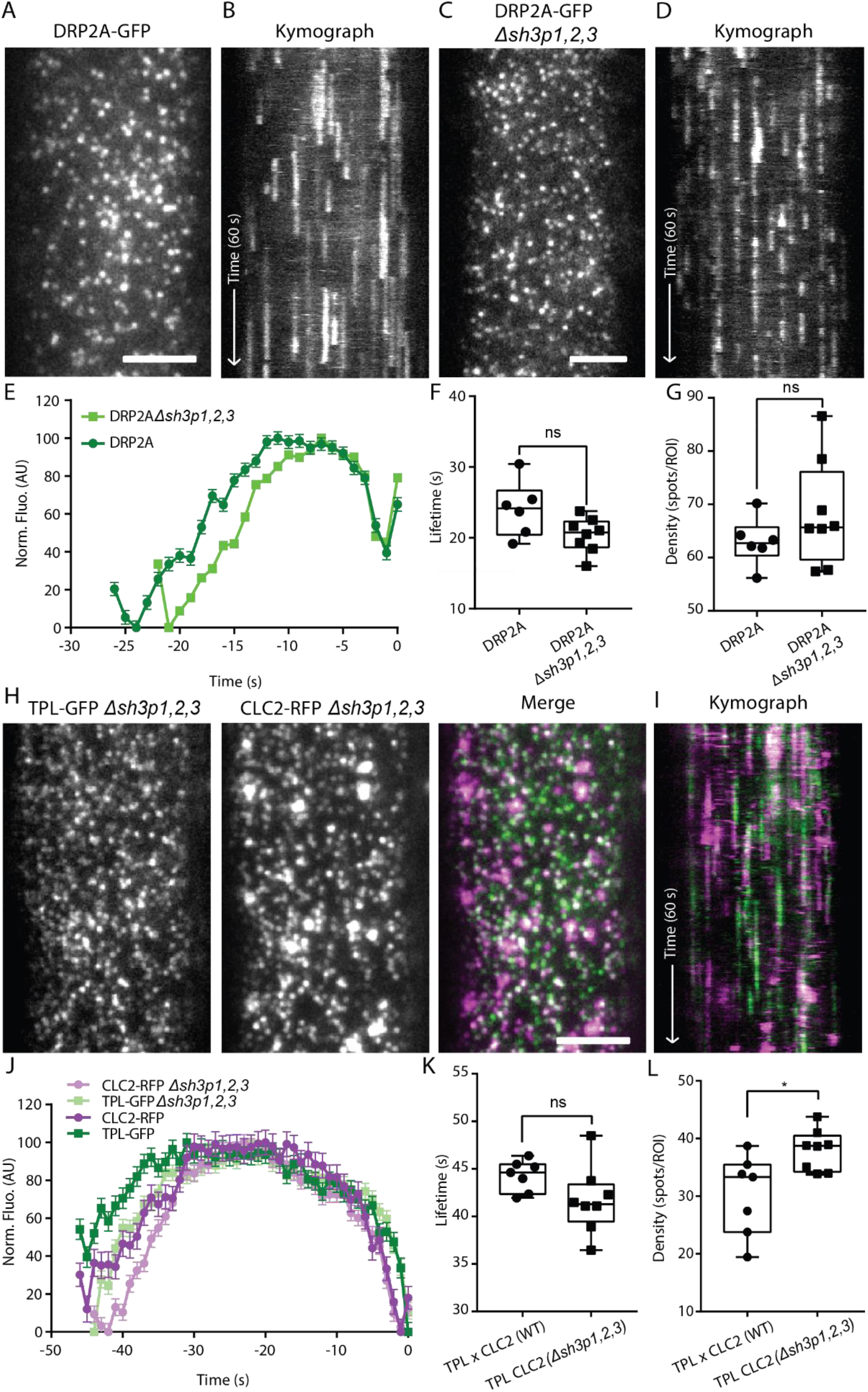
CME markers have a normal lifetime in *Δsh3p1,2,3* triple mutant. (A) TIRF-M images of a cell surface of root epidermal cell expressing DRP2A-GFP. Scale bar: 5 µm. (B) Representative kymograph of DRP2A lifetimes on the PM. (C) TIRF-M images of a cell surface of root epidermal cell expressing DRP2A-GFP in *Δsh3p1,2,3* background. Scale bar: 5 µm. (D) Representative kymograph of DRP2A in *Δsh3p1,2,3* lifetimes on the PM. (E-G) Data from six independent experiments for DRP2A and eight independent experiments for DRP2A in *Δsh3p1,2,3* were combined to generate a (E) mean recruitment profile of DRP2A foci, (F) mean lifetime (DRP2A, 24.45±0.1 s; DRP2A *Δsh3p1,2,3* 20.1±0.08 s), and (G) mean density (DRP2A, 62.97±1.8 spots ROI^−1^; DRP2A *Δsh3p1,2,3* 68.24±3.5 spots ROI^−1^) of CME events. Plots indicate Mean±SEM, DRP2A, n=6 cells from independent roots, 41,473 tracks; DRP2A *Δsh3p1,2,3*, n=8 cells from independent roots, 77,361 tracks. Plot, Mean ± SEM, ns> 0.05, t-test to compare to control. Scale bar: 5 µm. (H) TIRF-M images of a cell surface of root epidermal cell expressing TPL-GFP x CLC2-tagRFP in *Δsh3p1,2,3* background. Scale bar: 5 µm. (I) Representative kymograph of TPL-GFP x CLC2-tagRFP in *Δsh3p1,2,3* background lifetimes on the PM. (J-L) Data from seven independent experiments for TPL-GFP x CLC2-tagRFP and eight independent experiments for TPL-GFP x CLC2-tagRFP in *Δsh3p1,2,3* were combined to generate a (J) mean recruitment profile of TPL-GFP x CLC2-tagRFP and TPL-GFP x CLC2-tagRFP in *Δsh3p1,2,3* foci, (K) mean lifetime (TPL-GFP x CLC2-tagRFP, 44.2±0.3 s; TPL-GFP x CLC2-tagRFP *Δsh3p1,2,3,* 41.54±0.2 s), and (L) mean density (TPL-GFP x CLC2-tagRFP, 30.29±2.6 spots ROI^−1^; TPL-GFP x CLC2-tagRFP *Δsh3p1,2,3,* 38.02±1.2 spots ROI^−1^) of CME events. Plots indicate Mean±SEM, TPL-GFP x CLC2-tagRFP, n=7 cells from independent roots, 16,367 tracks; TPL-GFP x CLC2-tagRFP *Δsh3p1,2,3*, n=8 cells from independent roots, 24,217 tracks. Plot, Mean ± SEM, *P < 0.0155; ns> 0.05, t-test to compare to control.

In summary, the impaired membrane uptake observed in the Δ*sh3p1,2,3* triple mutant was not caused by changes in functioning of CME components, like DRP2A CLC or TPL.

### PIN2 recycling is not impaired in Δ*sh3p1,2,3* triple mutant

Previous reports show the SH3P2 binds ubiquitinated cargo and participates in vesicle trafficking (Nagel et al., 2017; Lam et al., 2001). Therefore, while SH3P2 might not have a major function during the process of CCV formation, it could potentially bind cargo immediately after vesicle has been cut and clathrin coat has been disassembled, thus explaining why we detect no change in the dynamics of CME markers at the PM in D*sh3p1,2,3* mutant.

To test this hypothesis, we analysed the recycling of PIN2 in D*sh3p1,2,3* mutant, as this protein undergoes K63 ubiquitination and constant recycling on PM through CME (Feraru et al., 2012; Leitner et al., 2012). Its recycling and polarity is dependent on proper functioning of CME machinery (Kitakura et al., 2011; Mravec et al., 2011). We used immunostaining of PIN2 in *A. thaliana* roots treated with Brefeldin A (BFA), which blocks protein recycling at Golgi apparatus (Naramoto et al., 2014). To observe the dynamics of only the PIN2 that was internalised through endocytosis and not synthesised *de novo* we pre-treated samples with Cycloheximide (CHX) (Schneider-Poetsch et al., 2010). An impairment in CME or trafficking of PIN proteins in D*sh3p1,2,3* mutant would result in reduced amounts and/or size of BFA bodies. Surprisingly, analysis of PIN2 immunostaining showed only minor difference in amount and size of BFA formations compared to the control (Fig. 7, S10), suggesting that in D*sh3p1,2,3* mutant PIN2 has normal internalisation and recycling rates.

**Fig. 7.**
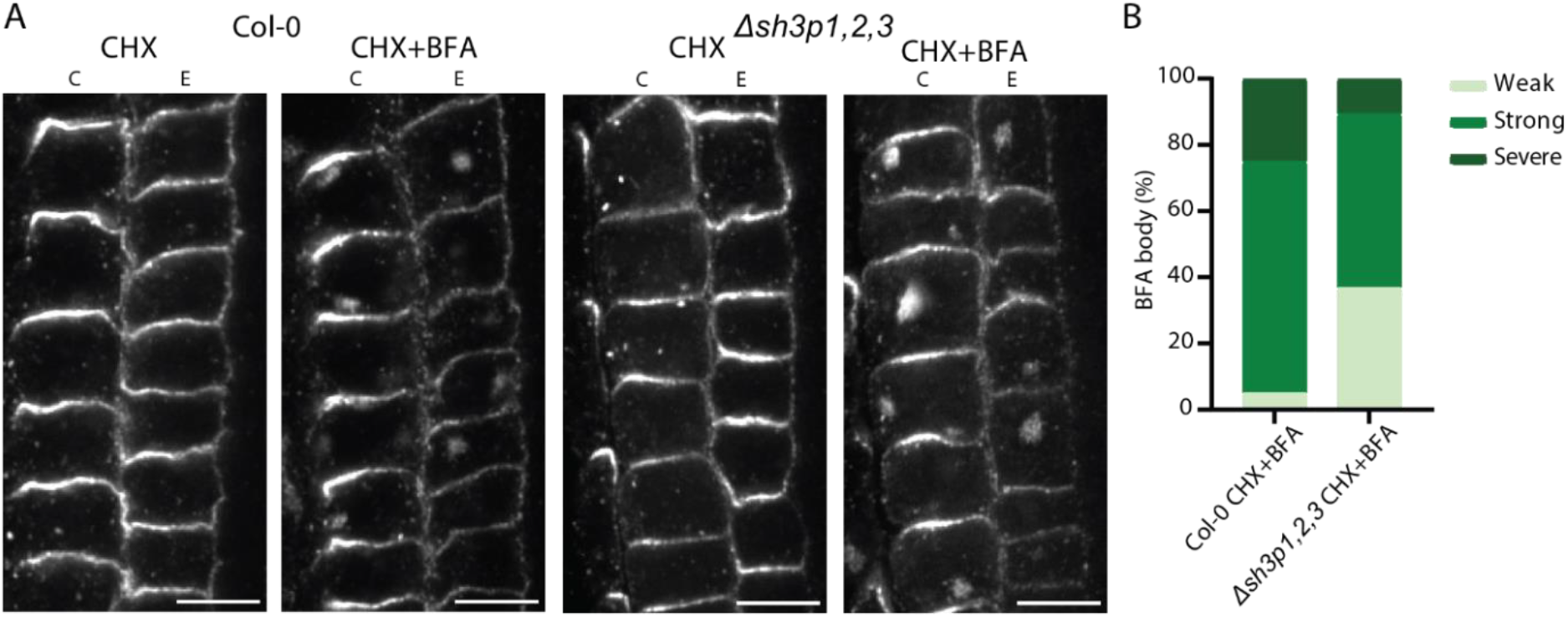
Recycling of PIN2 proteins is normal in *Δsh3p1,2,3* triple mutant. (A) Confocal images of PIN2 localization in Epidermis E and Cortex C cells of Arabidopsis roots treated with CHX alone (50 µM for 1h) or with BFA and CHX together (50 µM for 1h) in Col-0 and in Δsh3p1,2,3 triple mutant. (B) A stacked column chart representing the percentage of roots, per genotype, distributed in three different categories (weak, strong, severe) based on the abundance of PIN2 BFA bodies (internalization) formed in the cells after BFA treatment. Scale bar 20 µm.

These observations are puzzling, as despite the sever phenotype and reduced levels of membrane internalisation observed with FM4-64, CME rates and cargo internalisation in D*sh3p1,2,3* mutant plants are not significantly affected. This raises more questions about endocytosis in plants and function of SH3P proteins at the end of CCV formation.

## Discussion

The scission machinery of CCV is arguably one of the most important protein complexes in the CME mechanism. While the scission process is well-characterized in mammalian and yeast systems, there are significant gaps in our understanding of how CCVs are released from the plant’s PM. Members of the DRP2 family, DRP2A and DRP2B, play crucial role in plant growth and development (Backues et al., 2010). They have been hypothesised to do so through performing a similar scission function as their dynamin homologue in mammalian systems. Although some studies have shown co-localisation of DRP2s with other CME markers on PM, it has not yet been clear when and how they arrive at the site of CCV formation. In this study, we aimed to gain insight into the composition of the potential scission machinery and its recruitment mechanism during plant CME. Specifically, we wanted to determine when DRP2s are recruited during the process of CCV formation and whether SH3Ps play an important role in this recruitment.

Using high-resolution live imaging of DRP2s and SH3P2 together with CLC2 on the PM of root epidermal cells allowed us to study the dynamics of these proteins in the scope of CCV formation, maturation and removal from the membrane. We found that both DRP2s, as well as SH3P2, arrive close to the departure of CCV in the majority of endocytic events. Not only they arrive and co-localize during endocytosis, but they interact as confirmed by YTH and pull-down assays. Using an *in vitro* approach, we established the specific motif within DRP2s interacting with SH3 domain of SH3P2. Moreover, our *in vitro* data show that SH3P2 can deform membranes. These results support previously published data for DRP2s and SH3P2, and are also in line with the recruitment dynamics and biochemical properties of its mammalian and yeast homologues. However, our *in vivo* experiments with DRP2-PRM2 mutants, that could not interact with SH3P2 in YTH assays, showed no difference in DRP2-PRM2 nor CLC2 dynamics. What could be an explanation for this phenomenon? Closer analysis of recruitment profiles and co-localisation of DRP2A and SH3P2 suggests, that a) recruitment of DRP2A starts before the recruitment of SH3P2, and b) a significant amount of DRP2A is not co-localising with SH3P2, suggesting, that DRP2s can be recruited through other mechanisms that do not involve SH3P2. Potentially, other members of DRP family are able to interact and recruit each other, like DRP1s or DRP2B, as GTPases can polymerase through GED domain rather than PRD, and therefore recruit DRP2s with mutated PRM2 (Chappie et al., 2010; Fujimoto et al., 2008).

Next, we further investigated the functions of SH3P2 in CME. We used a recently generated triple mutant line that has a reduction of SH3P protein levels, to study endocytosis rates (Adamowski et al., 2022). Contradictory to the decrease in overall membrane internalization in D*sh3p1,2,3*, we observed normal dynamics and density of DRP2A, CLC2 as well as TPLATE CME markers. The analysis of PIN2 dynamics in D*sh3p1,2,3* mutant also did not show any changes compared to the control, indicating normal cargo recruitment and internalisation during CME. It needs to be mentioned, that FM uptake has not always been a reliable representation of endocytosis in plants. An earlier study shows that although plants with a mutation in clathrin heavy chain have significantly reduced FM uptake, the cargo recycling in these plants is not affected (Kitakura et al., 2011). These unexpected results raise further questions about the SH3P2 function in CME and the unexplained behaviour of D*sh3p1,2,3* and show the need for further studies of these proteins.

While our research was focused on the role of DRP2s and SH3P2 in plant CME, these proteins can be potentially involved in other types of endocytosis that are independent of clathrin. However, our knowledge about the plant Clathrin-Independent Endocytosis (CIE) and proteins mediating it is very limited. Interestingly, the N-BAR domain of SH3P2 is homologous to that of the mammalian Endophilin which, among its functions in CME, mediates a Dynamin-dependant CIE pathway called Fast Endophilin Mediated Endocytosis (FEME) (Ahn et al., 2017; Casamento and Boucrot, 2020). FEME pathway serves as a main internalisation road for fast (5 – 10 s) cargo internalisation, like the β1-adrenergic receptor (β1AR) and cytokine receptors (Boucrot et al., 2015; Ferreira et al., 2020). In plants, partially characterised CIE suggests the involvement of Flot1, a homologue to mammalian flotillin that mediates membrane bending during CIE, which is a Dynamin-dependant process (Glebov et al., 2006; Li et al., 2012). Taking into account that CLC2 foci have a higher co-localisation rate with SH3P2 and DRP2s than their co-localization rate with clathrin, it gives an indication of clathrin-independent DRP2-dependant endocytosis pathway in plants. Moreover, the ability of SH3P2 to deform membranes, partial co-localisation of SH3P2 with clathrin foci, and the lack of effects of SH3Ps mutation on the CME dynamics, it could be speculated that SH3P2 can play a role in yet-to-be-discovered CIE pathway; however, more research needs to be dedicated to discovering the exact mechanism of such potential pathway and cargoes, that can be internalised through it.

In conclusion, our study provides detailed insights into the dynamics and interaction between DRP2 and SH3P2 proteins during CME in plants. We found that while DRP2s and SH3P2 arrive at the end of CCV formation and show specific interaction *in vitro*, DRP2 is likely to be recruited through other, yet-to-be-uncovered mechanisms, rather than exclusively through SH3P2. These results provide further evidence that although plants possess many homologues of the mammalian and yeast CME system, plant CME functions in a rather unique and often unpredictable way.

## Material and Methods

### Plant material and growth conditions

All *Arabidopsis thaliana* mutants and transgenic lines used are in Columbia-0 (Col-0) background. Arabidopsis thaliana accession codes for genes used in this study: DRP2A (AT1G10290), DRP2B (AT1G59610), SH3P2 (AT4G34660.1), CLC2 (AT2G40060), TPLATE (AT3G01780), PIN1 (AT1G73590), PIN2 (AT5G57090). Transgenic Arabidopsis thaliana plants used in this study were pDRP2A::DRP2A-GFP *(drp2a-/-),* pDRP2A::DRP2A-GFP *(drp2a-/-;drp2b-/-),* pDRP2A::DRP2A-GFP × pRPS5A::CLC2-tagRFP, pDRP2A::DRP2A-PRM2-GFP × pRPS5A::CLC2-tagRFP, pDRP2B::DRP2B-GFP × pRPS5A::CLC2-tagRFP, pDRP2B::DRP2B-PRM2-GFP × pRPS5A::CLC2-tagRFP, pSH3P2::SH3P2-GFP × pRPS5A::CLC2-mOrange, pDRP2A::DRP2A-GFP × pSH3P2::SH3P2-tagRFP, *Δsh3p1,2, Δsh3p3, Δsh3p1,2,3,* pXVE:UBQ10::mCherry × pRPS5A::CLC2-GFP, pXVE:UBQ10::SH3-mCherry × pRPS5A::CLC2-GFP, pDRP2A::DRP2A-GFP, pDRP2A::DRP2A-GFP *(Δsh3p1,2,3),* pSH3P2::SH3P2-GFP, pSH3P2::SH3P2-GFP *(Δsh3p1,2,3),* pRPS5A::CLC2-GFP, pRPS5A::CLC2-GFP *(Δsh3p1,2,3),* pLAT52p::TPLATE-GFP × pRPS5A::CLC2-tagRFP, pLAT52p::TPLATE-GFP × pRPS5A::CLC2-tagRFP *(Δsh3p1,2,3)* (Adamowski et al., 2022; Backues et al., 2010a; Gadeyne et al., 2014b; Smith et al., 2014)*. Arabidopsis thaliana* seeds were surface-sterilized by either chlorine gas or 99% ethanol for 10 min. Seeds were then plated onto 1/2-Murashige-Skoog (MS) agar plates with 1% (weight/volume) sucrose, stratified for 1 to 2 d in the dark at 4 °C, and then transferred to the growth room (21 °C, 16 h light, 8 h dark) and grown vertically for 5 or 7 d depending on the type of experiment for which they were required. Light sources used were Philips GreenPower LED production modules [in deep red (660 nm)/far red (720 nm)/blue (455 nm) combination, Philips], with a photon density of 140.4 μmol/m2/s ± 3%.

### Cloning

Constructs for generating plant fluorescent reporter lines were generated using the Gate Way protocol (Thermo Fisher Scientific). Shortly, promoter sequences were amplified from gDNA generated from Col-0 plants and put in the p41r entry vector. Gene sequences were cloned using cDNA generated from Col-0 plants into pDONR221 entry vector. Fluorescent tags (GFP or mCherry) were cloned into p23 entry vector. The final plasmids were assembled using p41r, pDONR221 and p23 entry vectors and destination vectors with desired antibiotic resistance. *Agrobacterium tumefaciens* strain pGV3101 was used to deliver destination constructs into corresponding plants by floral dip.

Cloning of GAD-SH3P2, MBP-SH3P2(FL) and GST-SH3P2(SH3) was previously described in Nagel. et al, 2017. For GAD-SH3P2(SH3) and GAD-SH3P2(BAR), the fragments of SH3P2 were amplified with primers MN176-MS12 (SH3P2(SH3)) and MS11-MN177 (SH3P2(BAR)) and cloned between the EcoRI and XhoI and EcoRI and BamHI restriction sites of pGADT7 (Clontech) respectively. To obtain GBD-DRP1A(C406), GBD-DRP2A(C700) and GBD-DRP2B(C700) the fragments were amplified from Arabidopsis cDNA with the primer pairs MN349-MN226 (DRP1A(C406)), MN316-MN321 (DRP2A(C700)), and MN317-MN315 (DRP2B(C700)) and cloned between the NdeI and XhoI (DRP1A(C406)) and NdeI and EcoRI(DRP2A/B(C700)) restriction sites of pGBKT7. Mutated versions were amplified from DRP2B pDONR221 construct used as template using primer pairs MN317-MN315 and cloned between NdeI and EcoRI (DRP2A/B(C700)) restriction sites of pGBKT7. GBD-DRP2B(PRD1) and GBD-DRP2B(PRD2) were amplified with primers MN317-MN350 (DRP2B(PRD1)) and MN351-MN315 DRP2B(PRD2) and cloned between NdeI and EcoRI restriction sites of pGBKT7. GST-DRP2B(C474) was amplified by primer pairs MN256-MN255 and cloned between the EcoRI and XhoI restriction sites of pGEX-6p-1.

Constructs for protein overexpression in bacteria were generated using a Gibson Assembly protocol. GBlock gene with overhangs for the destination vector sequence was codon-optimized for *Escherichia coli* expression using the IDT service and then inserted into a pTB146 destination vector. A His-SUMO tag was located on the N-terminus of the expression construct. A list of primers used for cloning is listed in Supplementary Table 1.

### FM uptake assay, TIRF-M Imaging and Analysis

A Zeiss inverted LSM-800 confocal microscope equipped with Airyscan and 40× water immersion objected was used to examine the FM4-64 uptake in 7-d-old seedlings. Seedlings were incubated for 15 min at room temperature with 2 µM FM4-64 in AM+ media, washed three times in AM+ media, and imaged and analyzed, as described previously (Johnson et al., 2020). For TIRF-M experiments an Olympus IX83 inverted microscope equipped with a Cell^TIRF module using an OLYMPUS Uapo N 100×/1.49 Oil TIRF objective was used. Cut roots were mounted on the slide in AM+ media and covered with Precision Cover Glasses (refractive index 1.5H). Root epidermal cells were imaged and data was analyzed as described previously (Johnson et al., 2020) . Co-localization of proteins *in vivo* was done using Spots co-localization (ComDet v.0.5.5) plugin for ImageJ (https://github.com/UU-cellbiology/ComDet).

### YTH Assay

GAD- and GBD-fusion constructs were co-transformed into yeast strain Y8800 (based on PJ69-4; MATa, leu2-3,112 trp1-901 his3-200 ura3-52 gal4Δ gal80Δ GAL2-ADE2 LYS2::GAL1-HIS3 MET2::GAL7-lacZ cyh2R). Transformants were selected after 3 d on synthetic complete (SC) medium lacking leucine and tryptophan (−LW) at 30 °C. To examine reporter gene expression, transformants were grown on solid SC medium lacking leucine, tryptophan, and histidine (−LWH) supplemented with 5 mM 3-amino-1,2,4-triazole for 2 d at 30 °C. Yeast total proteins were extracted as described previously (67), and expression of constructs was analysed by immunoblotting using anti-GBD and anti-HA (GAD) antibodies.

### Protein Purification and *in vitro* Binding Assays

GST- or MBP-fused proteins were expressed in the Escherichia coli Rosetta (DE3), Rosetta-gami 2, or Rosetta-gami B strains (all from Merck Millipore) and purified using Pierce Glutathione Magnetic Beads (Thermo Scientific), or Amylose Resin (New England Biolabs), depending on the tag of the fusion protein. For SH3P2(SH3) the GST tag was removed by PreScission Protease (Cytiva). For in vitro-binding assays, Pierce Glutathione Magnetic Beads saturated with 80 pmol of GST-fusion proteins GST-DRP2B(C747) were incubated with an equimolar amount of MBP-SH3P2 or untagged SH3P2(SH3) in 400 μL of cold buffer (50 mM Tris·HCl at pH 7.5, 150 mM NaCl, 10 mM MgCl2, 0.05% Tween-20) under rotation at 4 °C. The beads were then washed four times with cold buffer, proteins were eluted with 40 mM glutathione. Bead bound materials were subjected to SDS/PAGE and analyzed by immunoblotting.

### Purification of SH3P2 protein

SH3P2 was cloned into vector pTB146, with an N-terminal 6xHis tag followed by SUMO fusion protein using Gibson assembly protocol (Thermo Fisher Scientific). Protein was expressed in *E. coli* BL21 cells, grown at 30 °C (250rpm) in LB medium supplemented with 100 µg/ml^−1^ ampicillin. Prior to expression the cell culture was cooled down to 4°C, and expression was induced at an OD600 of 0.8 with 1 mM IPTG. The protein was expressed overnight at 12 °C and harvested by centrifugation (5000 g for 30 min at 4 °C). The pellet was resuspended in lysis buffer (50 mM Tris-HCl pH8.0, 400 mM NaCl, 25 mM KH_2_PO_4_ pH8.0, 10% Glycerol, 5 mM EDTA, 5 mM DTT, 1% Triton X-100, 1 mM PMSF) supplemented with EDTA-free protease inhibitor cocktail tablets and 1 mg/ml^−1^ DNase I and 1 mg/ml^−1^ lysozyme. Cells were lysed by sonication using a Q700 Sonicator equipped with a probe of 12.7 mm diameter, which was immersed into the resuspended pellet. The suspension was kept on ice during sonication (Amplitude 25, 1 s ON and 4 s OFF for a total time of 10 min). Subsequently, cell debris was removed by centrifugation at 60,000 g for 1 h at 4 °C. The clarified lysate was incubated with Ni-NTA resin for 1 h at 4 °C. Subsequently, the resin was washed with 20x CV washing buffer A (50 mM Tris-HCl pH8.0, 400mM NaCl, 25mM KH_2_PO_4_ pH8.0, 10% Glycerol, 5mM EDTA, 5mM DTT, 0.2% Triton X-100) and 40x CV washing buffer B (50 mM Tris-HCl pH8.0, 400 mM NaCl, 25 mM KH_2_PO_4_ pH8.0, 10% Glycerol, 5 mM EDTA, 5 mM DTT, 20 mM Imidazole). Fusion protein was eluted using elution buffer (50 mM Tris-HCl pH8.0, 400 mM NaCl, 25 mM KH_2_PO_4_ pH8.0, 10% Glycerol, 5 mM EDTA, 5 mM DTT) containing increasing concentration of imidazole (50 - 300 mM). The protein concentration was determined with Bradford. The 6xHis tagged protease UlpI was added in a 1:100 molar ratio, and the 6xHis-SUMO tag was cleaved overnight at 4 °C, accompanied with dialysis in dialysis buffer (50 mM Tris-HCl pH8.0, 400 mM NaCl, 25 mM KH_2_PO_4_ pH8.0, 10% Glycerol, 5 mM EDTA, and 5 mM DTT) with gentle stirring. To exchange SH3P2 into the final buffer (25 mM HEPES pH7.5, 200 mM NaCl, 10% Glycerol, 1 mM EDTA, 5 mM DTT, and 25 mM KH_2_PO_4_ pH8.0) PD10 columns were used. To remove the cleaved tag and UlpI protease, SH3P2 was subjected to reverse affinity chromatography. The purity of the final protein was determined via SDS-page gel-electrophoresis and protein concentration was determined via NanoDrop and Bradford. Protein was flash-frozen in liquid N_2_ and stored at −80°C.

### LUV preparation

LUVs were prepared using a mixture of 1,2-dioleoyl-sn-glycero-3-phospho-(1’-rac-glycerol), 1,2-dioleoyl-sn-glycero-3-phospho-L-serine, 1,2-dioleoyl-sn-glycero-3-phosphate (at a ratio of 60:20:20 mol%) and 1,2-dioleoyl-sn-glycero-3-phospho-(1’-myo-inositol-4’,5′-bisphosphate) (PI(4,5)P_2_) (Avanti) at a ratio of DOPC (100, mol%), DOPC:DOPS (80:20, mol%), DOPC:DOPS:PA (80:18:2, mol%), DOPC:DOPS:PI(4,5)P_2_ (80:17.5:2,5 mol%). Lipids were mixed in a glass vial at the desired ratio, blow-dried with filtered N_2_ to form a thin homogeneous film, and kept under vacuum for 2 to 3 h. After, lipid film was rehydrated in a swelling buffer (25mM HEPES pH7.5, 200 mM NaCl, 25 mM KH_2_PO_4_ pH8.0) for 10 min at room temperature. Total lipid concentration was 2 mM. The mixture was vortexed rigorously, and the resulting dispersion of multilamellar vesicles was repeatedly freeze thawed (five to six times) in liquid N_2_. The mixture was extruded through a polycarbonate membrane with a pore size 400 nm (LiposoFast Liposome Factory). LUVs were stored at 4 °C and used within 4 d.

### Sedimentation assay

To assess membrane binding capacity of SH3P2 the pelleting assay was used. Shortly, LUVs were prepared as described above with swelling buffer containing 200 mM Sucrose instead of NaCl. SH3P2 protein was cleaned from aggregates using ultracentrifugation at 100 000 xg for 20 min at 4 °C. LUVs were incubated with the protein for 10 min at RT and spun down at 100 000 xg for 20 min at 4 °C. Supernatant was separated and pellet was suspended in outside buffer (25mM HEPES pH7.5, 200 mM NaCl, 25 mM KH_2_PO_4_ pH8.0). Samples were assessed SDS-page gel-electrophoresis. Images of gels were analyzed using GelAnalyzer 19.1 (www.gelanalyzer.com, by Istvan Lazar Jr., PhD and Istvan Lazar Sr., PhD, CSc).

### Tubulation Assay

To test membrane-bending activity of SH3P2, electron microscopy of LUVs incubated with 1 µM of SH3P2 was used. The final concentration of LUVs was 0.5 mM. Protein with LUVs or LUVs alone were incubated for 10 min at room temperature. A total of 20 μL mix was incubated on glow-discharged carbon-coated copper EM grids (300 mesh, EMS). Filter paper was used to remove any excess solution. Grids were then negatively stained with 2% uranyl acetate aqueous solution for 1 min and observed under a Tecnai 12 transmission electron microscope operated at 120 kV (Thermo Fisher Scientific). Images were analyzed using ImageJ.

### Mass photometer assay

To analyse the mass of individual protein molecules of SH3P2, a mass photometer assay was used. Coverslips were cleaned by sonication in mqH2O, Isopropanol and again mqH2O for at least 5 minutes each. Protein was diluted to 50 nM, 75 nM, and 100 nM final concentration in the final buffer (25 mM HEPES pH7.5, 200 mM NaCl, 25 mM KH_2_PO_4_ pH8.0). After adding the protein of interest, data was recorded at a framerate of 5ms/frame and recorded for 1 min. Results were analyzed using MP Discovery Software.

### Plant tissue immunostaining with BFA treatment

Whole-mount immunolocalization was performed on 4d old seedlings of Arabidopsis following the published protocol (Sauer and Friml, 2010). The seedlings were pre-treated with 50 µM Cycloheximide (CHX) for 30 minutes, followed by a co-treatment of CHX and Brefeldin A (BFA) for one hour with each at a concentration of 50 µM. The Antibodies rabbit anti-PIN2 (produced and processed in lab) were diluted 1:1000, and CY3-conjugated anti-rabbit secondary antibody (Sigma, C2306) 1:600. For confocal laser scanning microscopy, scans were taken using Zeiss LSM800.

## Acknowledgements

We thank prof. Eileen Lafer and Liping Wang for their suggestions regarding the optimization of protein expression and purification. We thank Maciek Adamowski for providing genetic material. This research was supported by the Scientific Service Units (SSU) of IST-Austria through resources provided by the Electron microscopy (EMF), Lab Support Facility (LSF) (particularly Dorota Jaworska) and the Bioimaging Facility (BIF).

## Author contributions

Conceptualization: N.G., A.J., M.L. and J.F.; Investigation: N.G., MK.N., A.M., A.J, A.H. material generation. Formal Analysis: N.G., A.J., MK.N., and A.M.; Writing – original draft: N.G., M.L. and J.F.; Writing – review & editing: all authors; Supervision: A.J. E.I., M.L., and J.F.

## Supplementary information

Supplementary information is available online at …

## References

Adamowski, M., Matijević, I., Narasimhan, M., Friml, J., 2022. SH3Ps recruit auxilin-like vesicle uncoating factors into clathrin-mediated endocytosis (preprint). Plant Biology. 10.1101/2022.01.07.475403

Ahn, G., Kim, H., Kim, D.H., Hanh, H., Yoon, Y., Singaram, I., Wijesinghe, K.J., Johnson, K.A., Zhuang, X., Liang, Z., Stahelin, R.V., Jiang, L., Cho, W., Kang, B.-H., Hwang, I., 2017. SH3 Domain-Containing Protein 2 Plays a Crucial Role at the Step of Membrane Tubulation during Cell Plate Formation. Plant Cell 29, 1388–1405. 10.1105/tpc.17.00108

Antonny, B., Burd, C., De Camilli, P., Chen, E., Daumke, O., Faelber, K., Ford, M., Frolov, V.A., Frost, A., Hinshaw, J.E., Kirchhausen, T., Kozlov, M.M., Lenz, M., Low, H.H., McMahon, H., Merrifield, C., Pollard, T.D., Robinson, P.J., Roux, A., Schmid, S., 2016. Membrane fission by dynamin: what we know and what we need to know. EMBO J. 35, 2270–2284. 10.15252/embj.201694613

Backues, S.K., Korasick, D.A., Heese, A., Bednarek, S.Y., 2010b. The *Arabidopsis* Dynamin-Related Protein2 Family Is Essential for Gametophyte Development. Plant Cell 22, 3218–3231. 10.1105/tpc.110.077727

Baquero Forero, A., Cvrčková, F., 2019. SH3Ps—Evolution and Diversity of a Family of Proteins Engaged in Plant Cytokinesis. Int. J. Mol. Sci. 20, 5623. 10.3390/ijms20225623

Barberon, M., Zelazny, E., Robert, S., Conéjéro, G., Curie, C., Friml, J., Vert, G., 2011. Monoubiquitin-dependent endocytosis of the IRON-REGULATED TRANSPORTER 1 (IRT1) transporter controls iron uptake in plants. Proc. Natl. Acad. Sci. 108. 10.1073/pnas.1100659108

Bashline, L., Li, S., Anderson, C.T., Lei, L., Gu, Y., 2013. The Endocytosis of Cellulose Synthase in Arabidopsis Is Dependent on μ2, a Clathrin-Mediated Endocytosis Adaptin. Plant Physiol. 163, 150–160. 10.1104/pp.113.221234

Bhatia, V.K., Madsen, K.L., Bolinger, P.-Y., Kunding, A., Hedegård, P., Gether, U., Stamou, D., 2009. Amphipathic motifs in BAR domains are essential for membrane curvature sensing. EMBO J. 28, 3303–3314. 10.1038/emboj.2009.261

Blood, P.D., Voth, G.A., 2006. Direct observation of Bin/amphiphysin/Rvs (BAR) domain-induced membrane curvature by means of molecular dynamics simulations. Proc. Natl. Acad. Sci. 103, 15068–15072. 10.1073/pnas.0603917103

Bolte, S., Talbot, C., Boutte, Y., Catrice, O., Read, N.D., Satiat-Jeunemaitre, B., 2004. FM-dyes as experimental probes for dissecting vesicle trafficking in living plant cells. J. Microsc. 214, 159–173. 10.1111/j.0022-2720.2004.01348.x

Boucrot, E., Ferreira, A.P.A., Almeida-Souza, L., Debard, S., Vallis, Y., Howard, G., Bertot, L., Sauvonnet, N., McMahon, H.T., 2015. Endophilin marks and controls a clathrin-independent endocytic pathway. Nature 517, 460–465. 10.1038/nature14067

Casamento, A., Boucrot, E., 2020. Molecular mechanism of Fast Endophilin-Mediated Endocytosis. Biochem. J. 477, 2327–2345. 10.1042/BCJ20190342

Chappie, J.S., Acharya, S., Leonard, M., Schmid, S.L., Dyda, F., 2010. G domain dimerization controls dynamin’s assembly-stimulated GTPase activity. Nature 465, 435–440. 10.1038/nature09032

Chen, X., Irani, N.G., Friml, J., 2011. Clathrin-mediated endocytosis: the gateway into plant cells. Curr. Opin. Plant Biol. 14, 674–682. 10.1016/j.pbi.2011.08.006

Claus, L.A.N., Savatin, D.V., Russinova, E., 2018. The crossroads of receptor-mediated signaling and endocytosis in plants: Endocytosis and signaling in plants. J. Integr. Plant Biol. 60, 827–840. 10.1111/jipb.12672

Dahhan, D.A., Reynolds, G.D., Cárdenas, J.J., Eeckhout, D., Johnson, A., Yperman, K., Kaufmann, W.A., Vang, N., Yan, X., Hwang, I., Heese, A., De Jaeger, G., Friml, J., Van Damme, D., Pan, J., Bednarek, S.Y., 2022. Proteomic characterization of isolated Arabidopsis clathrin-coated vesicles reveals evolutionarily conserved and plant-specific components. Plant Cell 34, 2150–2173. 10.1093/plcell/koac071

Dhonukshe, P., Aniento, F., Hwang, I., Robinson, D.G., Mravec, J., Stierhof, Y.-D., Friml, J., 2007. Clathrin-Mediated Constitutive Endocytosis of PIN Auxin Efflux Carriers in Arabidopsis. Curr. Biol. 17, 520–527. 10.1016/j.cub.2007.01.052

Di Rubbo, S., Irani, N.G., Kim, S.Y., Xu, Z.-Y., Gadeyne, A., Dejonghe, W., Vanhoutte, I., Persiau, G., Eeckhout, D., Simon, S., Song, K., Kleine-Vehn, J., Friml, J., De Jaeger, G., Van Damme, D., Hwang, I., Russinova, E., 2013. The Clathrin Adaptor Complex AP-2 Mediates Endocytosis of BRASSINOSTEROID INSENSITIVE1 in *Arabidopsis*. Plant Cell 25, 2986–2997. 10.1105/tpc.113.114058

Farsad, K., Ringstad, N., Takei, K., Floyd, S.R., Rose, K., De Camilli, P., 2001. Generation of high curvature membranes mediated by direct endophilin bilayer interactions. J. Cell Biol. 155, 193–200. 10.1083/jcb.200107075

Feraru, E., Feraru, M.I., Asaoka, R., Paciorek, T., De Rycke, R., Tanaka, H., Nakano, A., Friml, J., 2012. BEX5/RabA1b Regulates *trans*-Golgi Network-to-Plasma Membrane Protein Trafficking in *Arabidopsis*. Plant Cell 24, 3074–3086. 10.1105/tpc.112.098152

Ferreira, A.P.A., Casamento, A., Roas, S.C., Panambalana, J., Subramaniam, S., Schützenhofer, K., Halff, E.F., Wah Hak, L.C., McGourty, K., Kittler, J.T., Thalassinos, K., Martinvalet, D., Boucrot, E., 2020. Cdk5 and GSK3β inhibit Fast Endophilin-Mediated Endocytosis (preprint). Cell Biology. 10.1101/2020.04.11.036863

Fujimoto, M., Arimura, S., Nakazono, M., Tsutsumi, N., 2008. Arabidopsis dynamin-related protein DRP2B is co-localized with DRP1A on the leading edge of the forming cell plate. Plant Cell Rep. 27, 1581–1586. 10.1007/s00299-008-0583-0

Fujimoto, M., Arimura, S., Ueda, T., Takanashi, H., Hayashi, Y., Nakano, A., Tsutsumi, N., 2010. *Arabidopsis* dynamin-related proteins DRP2B and DRP1A participate together in clathrin-coated vesicle formation during endocytosis. Proc. Natl. Acad. Sci. 107, 6094–6099. 10.1073/pnas.0913562107

Gad, H., Ringstad, N., Löw, P., Kjaerulff, O., Gustafsson, J., Wenk, M., Di Paolo, G., Nemoto, Y., Crum, J., Ellisman, M.H., De Camilli, P., Shupliakov, O., Brodin, L., 2000. Fission and Uncoating of Synaptic Clathrin-Coated Vesicles Are Perturbed by Disruption of Interactions with the SH3 Domain of Endophilin. Neuron 27, 301–312. 10.1016/S0896-6273(00)00038-6

Gadeyne, A., Sánchez-Rodríguez, C., Vanneste, S., Di Rubbo, S., Zauber, H., Vanneste, K., Van Leene, J., De Winne, N., Eeckhout, D., Persiau, G., Van De Slijke, E., Cannoot, B., Vercruysse, L., Mayers, J.R., Adamowski, M., Kania, U., Ehrlich, M., Schweighofer, A., Ketelaar, T., Maere, S., Bednarek, S.Y., Friml, J., Gevaert, K., Witters, E., Russinova, E., Persson, S., De Jaeger, G., Van Damme, D., 2014. The TPLATE Adaptor Complex Drives Clathrin-Mediated Endocytosis in Plants. Cell 156, 691–704. 10.1016/j.cell.2014.01.039

Gallop, J.L., Jao, C.C., Kent, H.M., Butler, P.J.G., Evans, P.R., Langen, R., McMahon, H.T., 2006. Mechanism of endophilin N-BAR domain-mediated membrane curvature. EMBO J. 25, 2898–2910. 10.1038/sj.emboj.7601174

Glebov, O.O., Bright, N.A., Nichols, B.J., 2006. Flotillin-1 defines a clathrin-independent endocytic pathway in mammalian cells. Nat. Cell Biol. 8, 46–54. 10.1038/ncb1342

Habermann, B., 2004. The BAR-domain family of proteins: a case of bending and binding?: The membrane bending and GTPase-binding functions of proteins from the BAR-domain family. EMBO Rep. 5, 250–255. 10.1038/sj.embor.7400105

Heidstra, R., Sabatini, S., 2014. Plant and animal stem cells: similar yet different. Nat. Rev. Mol. Cell Biol. 15, 301–312. 10.1038/nrm3790

Hong, Z., Bednarek, S.Y., Blumwald, E., Hwang, I., Jurgens, G., Menzel, D., Osteryoung, K.W., Raikhel, N.V., Shinozaki, K., Tsutsumi, N., Verma, D.P.S., 2003. A unified nomenclature for Arabidopsis dynamin-related large GTPases based on homology and possible functions. Plant Mol. Biol. 53, 261–265. 10.1023/b:plan.0000007000.29697.81

Irani, N.G., Di Rubbo, S., Mylle, E., Van Den Begin, J., Schneider-Pizoń, J., Hniliková, J., Šíša, M., Buyst, D., Vilarrasa-Blasi, J., Szatmári, A.-M., Van Damme, D., Mishev, K., Codreanu, M.-C., Kohout, L., Strnad, M., Caño-Delgado, A.I., Friml, J., Madder, A., Russinova, E., 2012. Fluorescent castasterone reveals BRI1 signaling from the plasma membrane. Nat. Chem. Biol. 8, 583–589. 10.1038/nchembio.958

Jelínková, A., Malínská, K., Petrášek, J., 2019. Using FM Dyes to Study Endomembranes and Their Dynamics in Plants and Cell Suspensions, in: Cvrčková, F., Žárský, V. (Eds.), Plant Cell Morphogenesis, Methods in Molecular Biology. Springer New York, New York, NY, pp. 173–187. 10.1007/978-1-4939-9469-4_11

Jhaveri, A., Maisuria, D., Varga, M., Mohammadyani, D., Johnson, M.E., 2021. Thermodynamics and Free Energy Landscape of BAR-Domain Dimerization from Molecular Simulations. J. Phys. Chem. B 125, 3739–3751. 10.1021/acs.jpcb.0c10992

Jockusch, W.J., Praefcke, G.J.K., McMahon, H.T., Lagnado, L., 2005. Clathrin-Dependent and Clathrin-Independent Retrieval of Synaptic Vesicles in Retinal Bipolar Cells. Neuron 46, 869–878. 10.1016/j.neuron.2005.05.004

Johnson, A., Dahhan, D.A., Gnyliukh, N., Kaufmann, W.A., Zheden, V., Costanzo, T., Mahou, P., Hrtyan, M., Wang, J., Aguilera-Servin, J., Van Damme, D., Beaurepaire, E., Loose, M., Bednarek, S.Y., Friml, J., 2021. The TPLATE complex mediates membrane bending during plant clathrin–mediated endocytosis. Proc. Natl. Acad. Sci. 118, e2113046118. 10.1073/pnas.2113046118

Johnson, A., Gnyliukh, N., Kaufmann, W.A., Narasimhan, M., Vert, G., Bednarek, S.Y., Friml, J., 2020. Experimental toolbox for quantitative evaluation of clathrin-mediated endocytosis in the plant model *Arabidopsis*. J. Cell Sci. jcs.248062. 10.1242/jcs.248062

Kitakura, S., Vanneste, S., Robert, S., Löfke, C., Teichmann, T., Tanaka, H., Friml, J., 2011. Clathrin Mediates Endocytosis and Polar Distribution of PIN Auxin Transporters in *Arabidopsis*. Plant Cell 23, 1920–1931. 10.1105/tpc.111.083030

Kolb, C., Nagel, M.-K., Kalinowska, K., Hagmann, J., Ichikawa, M., Anzenberger, F., Alkofer, A., Sato, M.H., Braun, P., Isono, E., 2015. FYVE1 Is Essential for Vacuole Biogenesis and Intracellular Trafficking in Arabidopsis. Plant Physiol. 167, 1361–1373. 10.1104/pp.114.253377

Kontaxi, C., Cousin, M.A., 2023. The phospho-regulated amphiphysin/endophilin interaction is required for synaptic vesicle endocytosis (preprint). Neuroscience. 10.1101/2023.01.15.524101

Lam, B.C.-H., Sage, T.L., Bianchi, F., Blumwald, E., 2002. Regulation of ADL6 activity by its associated molecular network: Regulation of ADL6 by its interacting partners. Plant J. 31, 565–576. 10.1046/j.1365-313X.2002.01377.x

Lam, B.C.-H., Sage, T.L., Bianchi, F., Blumwald, E., 2001. Role of SH3 Domain–Containing Proteins in Clathrin-Mediated Vesicle Trafficking in Arabidopsis. Plant Cell 13, 2499– 2512. 10.1105/tpc.010279

Lebecq, A., Doumane, M., Fangain, A., Bayle, V., Leong, J.X., Rozier, F., Marques-Bueno, M.D., Armengot, L., Boisseau, R., Simon, M.L., Franz-Wachtel, M., Macek, B., Üstün, S., Jaillais, Y., Caillaud, M.-C., 2022. The Arabidopsis SAC9 enzyme is enriched in a cortical population of early endosomes and restricts PI(4,5)P2 at the plasma membrane. eLife 11, e73837. 10.7554/eLife.73837

Leitner, J., Petrášek, J., Tomanov, K., Retzer, K., Pařezová, M., Korbei, B., Bachmair, A., Zažímalová, E., Luschnig, C., 2012. Lysine ^63^-linked ubiquitylation of PIN2 auxin carrier protein governs hormonally controlled adaptation of *Arabidopsis* root growth. Proc. Natl. Acad. Sci. 109, 8322–8327. 10.1073/pnas.1200824109

Li, R., Liu, P., Wan, Y., Chen, T., Wang, Q., Mettbach, U., Baluška, F., Šamaj, J., Fang, X., Lucas, W.J., Lin, J., 2012. A Membrane Microdomain-Associated Protein, *Arabidopsis* Flot1, Is Involved in a Clathrin-Independent Endocytic Pathway and Is Required for Seedling Development. Plant Cell 24, 2105–2122. 10.1105/tpc.112.095695

Lu, R., Drubin, D.G., Sun, Y., 2016. Clathrin-mediated endocytosis in budding yeast at a glance. J. Cell Sci. 129, 1531–1536. 10.1242/jcs.182303

Luo, L., Xue, J., Kwan, A., Gamsjaeger, R., Wielens, J., Von Kleist, L., Cubeddu, L., Guo, Z., Stow, J.L., Parker, M.W., Mackay, J.P., Robinson, P.J., 2016. The Binding of Syndapin SH3 Domain to Dynamin Proline-rich Domain Involves Short and Long Distance Elements. J. Biol. Chem. 291, 9411–9424. 10.1074/jbc.M115.703108

Luschnig, C., Vert, G., 2014. The dynamics of plant plasma membrane proteins: PINs and beyond. Development 141, 2924–2938. 10.1242/dev.103424

Marks, B., Stowell, M.H.B., Vallis, Y., Mills, I.G., Gibson, A., Hopkins, C.R., McMahon, H.T., 2001. GTPase activity of dynamin and resulting conformation change are essential for endocytosis. Nature 410, 231–235. 10.1038/35065645

Martin, T.F.J., 2001. PI(4,5)P2 regulation of surface membrane traffic. Curr. Opin. Cell Biol. 13, 493–499. 10.1016/S0955-0674(00)00241-6

Mbengue, M., Bourdais, G., Gervasi, F., Beck, M., Zhou, J., Spallek, T., Bartels, S., Boller, T., Ueda, T., Kuhn, H., Robatzek, S., 2016. Clathrin-dependent endocytosis is required for immunity mediated by pattern recognition receptor kinases. Proc. Natl. Acad. Sci. 113, 11034–11039. 10.1073/pnas.1606004113

McMahon, H.T., Boucrot, E., 2011. Molecular mechanism and physiological functions of clathrin-mediated endocytosis. Nat. Rev. Mol. Cell Biol. 12, 517–533. 10.1038/nrm3151

Meinecke, M., Boucrot, E., Camdere, G., Hon, W.-C., Mittal, R., McMahon, H.T., 2013. Cooperative Recruitment of Dynamin and BIN/Amphiphysin/Rvs (BAR) Domain-containing Proteins Leads to GTP-dependent Membrane Scission*. J. Biol. Chem. 288, 6651–6661. 10.1074/jbc.M112.444869

Mravec, J., Petrášek, J., Li, N., Boeren, S., Karlova, R., Kitakura, S., Pařezová, M., Naramoto, S., Nodzyński, T., Dhonukshe, P., Bednarek, S.Y., Zažímalová, E., de Vries, S., Friml, J., 2011. Cell Plate Restricted Association of DRP1A and PIN Proteins Is Required for Cell Polarity Establishment in Arabidopsis. Curr. Biol. 21, 1055–1060. 10.1016/j.cub.2011.05.018

Nagel, M.-K., Kalinowska, K., Vogel, K., Reynolds, G.D., Wu, Z., Anzenberger, F., Ichikawa, M., Tsutsumi, C., Sato, M.H., Kuster, B., Bednarek, S.Y., Isono, E., 2017a. *Arabidopsis* SH3P2 is an ubiquitin-binding protein that functions together with ESCRT-I and the deubiquitylating enzyme AMSH3. Proc. Natl. Acad. Sci. 114. 10.1073/pnas.1710866114

Naramoto, S., Otegui, M.S., Kutsuna, N., De Rycke, R., Dainobu, T., Karampelias, M., Fujimoto, M., Feraru, E., Miki, D., Fukuda, H., Nakano, A., Friml, J., 2014. Insights into the Localization and Function of the Membrane Trafficking Regulator GNOM ARF-GEF at the Golgi Apparatus in *Arabidopsis*. Plant Cell 26, 3062–3076. 10.1105/tpc.114.125880

Narasimhan, M., Gallei, M., Tan, S., Johnson, A., Verstraeten, I., Li, L., Rodriguez, L., Han, H., Himschoot, E., Wang, R., Vanneste, S., Sánchez-Simarro, J., Aniento, F., Adamowski, M., Friml, J., 2021. Systematic analysis of specific and nonspecific auxin effects on endocytosis and trafficking. Plant Physiol. 186, 1122–1142. 10.1093/plphys/kiab134

Narasimhan, M., Johnson, A., Prizak, R., Kaufmann, W.A., Tan, S., Casillas-Pérez, B., Friml, J., 2020. Evolutionarily unique mechanistic framework of clathrin-mediated endocytosis in plants. eLife 9, e52067. 10.7554/eLife.52067

Okamoto, P.M., Herskovits, J.S., Vallee, R.B., 1997. Role of the Basic, Proline-rich Region of Dynamin in Src Homology 3 Domain Binding and Endocytosis. J. Biol. Chem. 272, 11629–11635. 10.1074/jbc.272.17.11629

Paciorek, T., Zažímalová, E., Ruthardt, N., Petrášek, J., Stierhof, Y.-D., Kleine-Vehn, J., Morris, D.A., Emans, N., Jürgens, G., Geldner, N., Friml, J., 2005. Auxin inhibits endocytosis and promotes its own efflux from cells. Nature 435, 1251–1256. 10.1038/nature03633

Pan, J., Fujioka, S., Peng, J., Chen, J., Li, G., Chen, R., 2009. The E3 Ubiquitin Ligase SCFTIR1/AFB and Membrane Sterols Play Key Roles in Auxin Regulation of Endocytosis, Recycling, and Plasma Membrane Accumulation of the Auxin Efflux Transporter PIN2 in *Arabidopsis thaliana*. Plant Cell 21, 568–580. 10.1105/tpc.108.061465

Pant, S., Sharma, M., Patel, K., Caplan, S., Carr, C.M., Grant, B.D., 2009. AMPH-1/Amphiphysin/Bin1 functions with RME-1/Ehd1 in endocytic recycling. Nat. Cell Biol. 11, 1399–1410. 10.1038/ncb1986

Perrais, D., 2022. Cellular and structural insight into dynamin function during endocytic vesicle formation: a tale of 50 years of investigation. Biosci. Rep. 42, BSR20211227. 10.1042/BSR20211227

Peter, B.J., Kent, H.M., Mills, I.G., Vallis, Y., Butler, P.J.G., Evans, P.R., McMahon, H.T., 2004. BAR Domains as Sensors of Membrane Curvature: The Amphiphysin BAR Structure. Science 303, 495–499. 10.1126/science.1092586

Postma, J., Liebrand, T.W.H., Bi, G., Evrard, A., Bye, R.R., Mbengue, M., Kuhn, H., Joosten, M.H.A.J., Robatzek, S., 2016. Avr4 promotes Cf-4 receptor-like protein association with the BAK1/SERK3 receptor-like kinase to initiate receptor endocytosis and plant immunity. New Phytol. 210, 627–642. 10.1111/nph.13802

Prichard, K.L., O’Brien, N.S., Murcia, S.R., Baker, J.R., McCluskey, A., 2022. Role of Clathrin and Dynamin in Clathrin Mediated Endocytosis/Synaptic Vesicle Recycling and Implications in Neurological Diseases. Front. Cell. Neurosci. 15, 754110. 10.3389/fncel.2021.754110

Renard, H.-F., Simunovic, M., Lemière, J., Boucrot, E., Garcia-Castillo, M.D., Arumugam, S., Chambon, V., Lamaze, C., Wunder, C., Kenworthy, A.K., Schmidt, A.A., McMahon, H.T., Sykes, C., Bassereau, P., Johannes, L., 2015. Endophilin-A2 functions in membrane scission in clathrin-independent endocytosis. Nature 517, 493–496. 10.1038/nature14064

Rooij, I.I.S., Allwood, E.G., Aghamohammadzadeh, S., Hettema, E.H., Goldberg, M.W., Ayscough, K.R., 2010. A role for the dynamin-like protein Vps1 during endocytosis in yeast. J. Cell Sci. 123, 3496–3506. 10.1242/jcs.070508

Rosendale, M., Van, T.N.N., Grillo-Bosch, D., Sposini, S., Claverie, L., Gauthereau, I., Claverol, S., Choquet, D., Sainlos, M., Perrais, D., 2019. Functional recruitment of dynamin requires multimeric interactions for efficient endocytosis. Nat. Commun. 10, 4462. 10.1038/s41467-019-12434-9

Sánchez-Rodríguez, C., Shi, Y., Kesten, C., Zhang, D., Sancho-Andrés, G., Ivakov, A., Lampugnani, E.R., Sklodowski, K., Fujimoto, M., Nakano, A., Bacic, A., Wallace, I.S., Ueda, T., Van Damme, D., Zhou, Y., Persson, S., 2018. The Cellulose Synthases Are Cargo of the TPLATE Adaptor Complex. Mol. Plant 11, 346–349. 10.1016/j.molp.2017.11.012

Sauer, M., Friml, J., 2010. Immunolocalization of Proteins in Plants, in: Hennig, L., Köhler, C. (Eds.), Plant Developmental Biology, Methods in Molecular Biology. Humana Press, Totowa, NJ, pp. 253–263. 10.1007/978-1-60761-765-5_17

Schmid, S.L., Frolov, V.A., 2011. Dynamin: functional design of a membrane fission catalyst. Annu. Rev. Cell Dev. Biol. 27, 79–105. 10.1146/annurev-cellbio-100109-104016

Schneider-Poetsch, T., Ju, J., Eyler, D.E., Dang, Y., Bhat, S., Merrick, W.C., Green, R., Shen, B., Liu, J.O., 2010. Inhibition of eukaryotic translation elongation by cycloheximide and lactimidomycin. Nat. Chem. Biol. 6, 209–217. 10.1038/nchembio.304

Shupliakov, O., Löw, P., Grabs, D., Gad, H., Chen, H., David, C., Takei, K., De Camilli, P., Brodin, L., 1997. Synaptic Vesicle Endocytosis Impaired by Disruption of Dynamin-SH3 Domain Interactions. Science 276, 259–263. 10.1126/science.276.5310.259

Smaczynska-de Rooij, I.I., Allwood, E.G., Mishra, R., Booth, W.I., Aghamohammadzadeh, S., Goldberg, M.W., Ayscough, K.R., 2012. Yeast Dynamin Vps1 and Amphiphysin Rvs167 Function Together During Endocytosis: Vps1 and Rvs167 Function Together in Endocytosis. Traffic 13, 317–328. 10.1111/j.1600-0854.2011.01311.x

Sundborger, A.C., Fang, S., Heymann, J.A., Ray, P., Chappie, J.S., Hinshaw, J.E., 2014. A Dynamin Mutant Defines a Superconstricted Prefission State. Cell Rep. 8, 734–742. 10.1016/j.celrep.2014.06.054

Szaszák, M., Gáborik, Z., Turu, G., McPherson, P.S., Clark, A.J.L., Catt, K.J., Hunyady, L., 2002. Role of the Proline-rich Domain of Dynamin-2 and Its Interactions with Src Homology 3 Domains during Endocytosis of the AT1 Angiotensin Receptor. J. Biol. Chem. 277, 21650–21656. 10.1074/jbc.M200778200

Taylor, M.J., Perrais, D., Merrifield, C.J., 2011a. A High Precision Survey of the Molecular Dynamics of Mammalian Clathrin-Mediated Endocytosis. PLoS Biol. 9, e1000604. 10.1371/journal.pbio.1000604

Trache, A., Meininger, G.A., 2008. Total Internal Reflection Fluorescence (TIRF) Microscopy: Microscopy. Curr. Protoc. Microbiol. 10. 10.1002/9780471729259.mc02a02s10

Wang, S., Yoshinari, A., Shimada, T., Hara-Nishimura, I., Mitani-Ueno, N., Feng Ma, J., Naito, S., Takano, J., 2017. Polar Localization of the NIP5;1 Boric Acid Channel Is Maintained by Endocytosis and Facilitates Boron Transport in Arabidopsis Roots. Plant Cell 29, 824–842. 10.1105/tpc.16.00825

Xin, X., Gfeller, D., Cheng, J., Tonikian, R., Sun, L., Guo, A., Lopez, L., Pavlenco, A., Akintobi, A., Zhang, Y., Rual, J., Currell, B., Seshagiri, S., Hao, T., Yang, X., Shen, Y.A., Salehi-Ashtiani, K., Li, J., Cheng, A.T., Bouamalay, D., Lugari, A., Hill, D.E., Grimes, M.L., Drubin, D.G., Grant, B.D., Vidal, M., Boone, C., Sidhu, S.S., Bader, G.D., 2013. SH3 interactome conserves general function over specific form. Mol. Syst. Biol. 9, 652. 10.1038/msb.2013.9

Yoshinari, A., Fujimoto, M., Ueda, T., Inada, N., Naito, S., Takano, J., 2016. DRP1-Dependent Endocytosis is Essential for Polar Localization and Boron-Induced Degradation of the Borate Transporter BOR1 in *Arabidopsis thaliana*. Plant Cell Physiol. 57, 1985–2000. 10.1093/pcp/pcw121

Youn, J.-Y., Friesen, H., Kishimoto, T., Henne, W.M., Kurat, C.F., Ye, W., Ceccarelli, D.F., Sicheri, F., Kohlwein, S.D., McMahon, H.T., Andrews, B.J., 2010. Dissecting BAR Domain Function in the Yeast Amphiphysins Rvs161 and Rvs167 during Endocytosis. Mol. Biol. Cell 21, 3054–3069. 10.1091/mbc.e10-03-0181

Young, G., Hundt, N., Cole, D., Fineberg, A., Andrecka, J., Tyler, A., Olerinyova, A., Ansari, A., Marklund, E.G., Collier, M.P., Chandler, S.A., Tkachenko, O., Allen, J., Crispin, M., Billington, N., Takagi, Y., Sellers, J.R., Eichmann, C., Selenko, P., Frey, L., Riek, R., Galpin, M.R., Struwe, W.B., Benesch, J.L.P., Kukura, P., 2018. Quantitative mass imaging of single biological macromolecules. Science 360, 423–427. 10.1126/science.aar5839

Yu, X., Cai, M., 2004. The yeast dynamin-related GTPase Vps1p functions in the organization of the actin cytoskeleton via interaction with Sla1p. J. Cell Sci. 117, 3839–3853. 10.1242/jcs.01239

Zhang, L., Xing, J., Lin, J., 2019. At the intersection of exocytosis and endocytosis in plants. New Phytol. 224, 1479–1489. 10.1111/nph.16018

Zhuang, X., Jiang, L., 2014. Autophagosome biogenesis in plants: Roles of SH3P2. Autophagy 10, 704–705. 10.4161/auto.28060

Zhuang, X., Wang, H., Lam, S.K., Gao, C., Wang, X., Cai, Y., Jiang, L., 2013. A BAR-Domain Protein SH3P2, Which Binds to Phosphatidylinositol 3-Phosphate and ATG8, Regulates Autophagosome Formation in Arabidopsis. Plant Cell 25, 4596–4615. 10.1105/tpc.113.118307

