## Supplementary data for "Role of Dynamin-Related Proteins 2 and SH3P2 in Clathrin-Mediated Endocytosis in Plants" (from another version of this preprint: version 1, 2023-10-10)

### Supplemental Data

| Primer name | Primer sequence |
|---|---|
| pTB146_fwd | TCGAGCCCGGGTGACTGCAGG |
| pTB146_rev | ACCACCAATCTGTTCTCTGTGAGCCTC |
| codOpt_SH3P2 | ATGGAGGATAACGATATTATTGAGGCTCACAGAGAACAGATTGGTGGTGATGCAAT<br>ACGGAAGCAGGCCTCGCGGCTGCGCGAGCAGGTAGCTAGACAGCAACAAGCTGTG<br>TTTAAACAGTTTGGAGGTGGGGGCTACGGCAGCGGCTTGGCAGACGAGGCTGAACT<br>GAACCAACATCAGAAACTTGAGAAGCTGTATATTTCTACACGGGCGGCAAAACACT<br>ATCAACGCGACATCGTTCGGGGCGTCGAAGGGTATATTGTTACTGGAAGCAAGCAA<br>GTCGAAATCGGTACAAAGCTGTCAGAAGACAGTCGTAAATACGGTAGTGAAAACAC<br>CTGTACAAACGGAAATGTGTTGACCCGTGCAGCGTTGAACTATGGTCGCGCCAGAG<br>CTCAAATGGAAAAGGAACGGGGTAATATGCTGAAAGCGCTTGGAACGCAAGTGGC<br>AGAACCTCTTCGGGCCATGGTGCTTGGGGCACCGTTGGAAGACGCGCGCCATCTGG<br>CTCAACGCTATGATCGCATGAGACAAGAAGCTGAAGCTCAGGCGACAGAGGTGGC<br>ACGTCGGCAGGCAAAGGCTCGTGAATCGCAAGGCAACCCGGACATATTGATGAAGT<br>TAGAATCTGCGGAAGCAAAACTTCACGATCTTAAAAGCAATATGACCATATTAGGA<br>AAGGAGGCAGCATCTGCACTGGCGTCAGTAGAAGACCAACAGCAAAAACTGACGT<br>TAGAACGTCTGTTGTCTATGGTGGAGTCCGAAAGAGCTTACCATCAGAGAGTTTTAC<br>AGATACTTGATCAGCTTGAGGGCGAAATGGTTTCAGAGCGTCAGCGCATTGAGGCG<br>CCATCCACACCCAGTTCAGCGGACTCGATGCCACCTCCTCCATCGTACGAGGAGGC<br>AAACGGAGTTTTCGCCAGTCAAATGCATGACACTTCTACCGACTCTATGGGATACTT<br>TCTGGGGGAGGTATTGTTCCCATACCACGGGGTAACAGATGTCGAATTGAGCTTATC<br>CACGGGTGAGTACGTTGTTGTGCGCAAGGTCACCGGTTCTGGATGGGCTGAAGGGG<br>AGTGTAAAGGGAAGGCGGGATGGTTTCCTTACGGGTACATTGAAAGACGTGAGCGT<br>GTATTGGCTTCAAAAGTTAGCGAGGTATTCTGATCGAGCCCGGGTGACTGCAGGAA<br>GGGGATCCGGCTGCTA |
| DRPR2A_prom_fw | GGGGACAACTTTGTATAGAAAAGTTGTTGTCCGCAGATCTTGCC |
| DRPR2A_prom_rev | GGGGACTGCTTTTTTGTACAAACTTGACGCGACTAGCAAAAGC |
| DRPR2B_prom_fw | GGGGACTGCTTTTTTGTACAAACTTGActtcttccactgtacacact |
| DRPR2B_prom_rev | GGGGACAACTTTTCTATACAAAGTTGTCtgttttgccgttcaaagaga |
| Drp2aGDNAAttB1 | GGGGACAAGTTTGTACAAAAAAGCAGGCTTCATGGAGGCGATCGATGAGTT |
| Drp2aGDNAAttB2 | GGGGACCACTTTGTACAAGAAAGCTGGGTAATACCTATAAGCTGAACCTG |
| Drp2bGDNAAttB1 | GGGGACAAGTTTGTACAAAAAAGCAGGCTTCATGGAGGCGATCGATGAGTT |
| Drp2bGDNAAttB2 | GGGGACCACTTTGTACAAGAAAGCTGGGTACTAATACCTGTAAGATGATC |
| Drp2a p902 903a F | gcggtggtgctgctgccaaccgattggg |
| Drp2a p902 903a R | cccaatcggttggcagcagcaccaccgc |
| Drp2b p908 909a F | gtggcggagctgctgcgagccggtttgg |
| Drp2b p908 909a F | ccaaaccggctcgcagcagctccgccac |
| SH3_dom_SH3P2_fwd | GGGGACAAGTTTGTACAAAAAAGCAGGCTTTATGGCATCTCAGATGCATGACA |
| SH3_dom_SH3P2_rev | GGGGACCACTTTGTACAAGAAAGCTGGGTAGAAAACTTCGGACACTTTGC |
| MN176 SH3P2(C257-369)fw EcoRI | AAGGGAATTCATGTCTGAGAGGCAACGTATAG |

| | |
|---|---|
| MN177 SH3P2(N1-257)rv BamHI | AAGGGGATCCTCATACCATCTCTCCTTCGAG |
| MN226 AT5G42080 XhoI rv | AAGGCTCGAGTCACTTGGACCAAGCA |
| MN255 DRP2A rv XhoI | AAGGCTCGAGCTAATACCTATAAGCTGAACCTGTAG |
| MN256 DRP2B(aa747) fw EcoRI | AAGGGAATTCTATGTTGAAGCTGTTCTCAAC |
| MN315 DRP2B (EcoRI) rv | AAGGGAATTCCTAATACCTGTAAGATGATCCAG |
| MN316 DRP2A(C700) fw (NdeI) | AAGGCATATGATGGGCCAAGTGGGCAGTG |
| MN317 DRP2B(C700) fw (NdeI) | AAGGCATATGATGAAGGTTATCCAGGCCC |
| MN321 DRP2A rv (EcoRI) | AAGGGAATTCCTAATACCTATAAGCTGAACCTGTAG |
| MN349 DRP1A (NdeI) fw | AAGGCATATGGGTTACCGTCGTCTCATTGA |
| MN350 DRP2B (700-800) rv (EcoRI) | AAGGGAATTCCTAGTTCTGATCCTCTTGAATCAATG |
| MN351 DRP2B(C837) fw (NdeI) | AAGGCATATGATGGACAATAGTGGCACTGAAAG |
| MS11 SH3P2 fw EcoRI | GGAAGAATTCAGTGATGCAATTAGAAAACA |
| MS12 SH3P2 rv SalI | GGAAGTCGACTCAGAAAACTTCGGACACT |

**Table 1 – Codon optimized AtSH3P2 and primers used**

### Supplemental Figures

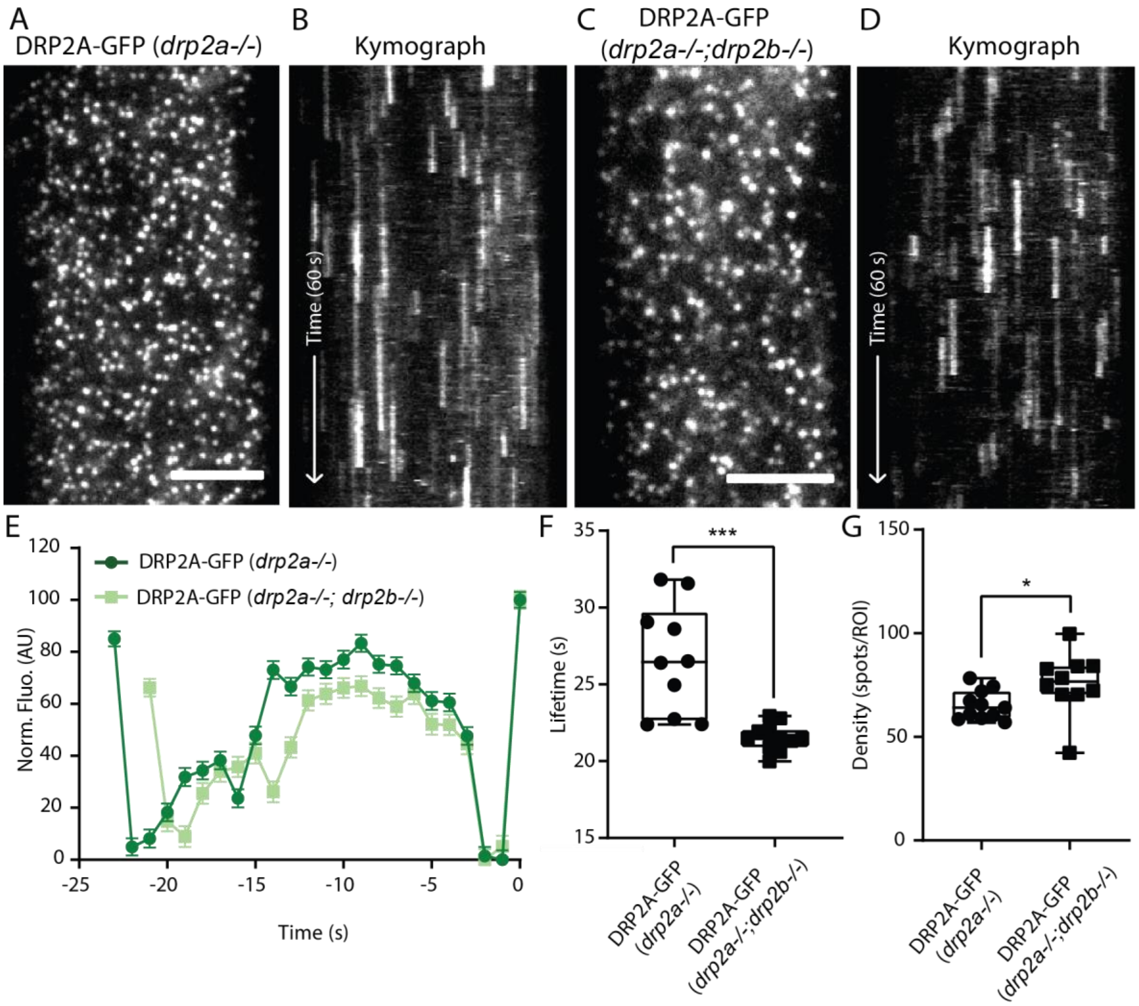


**Fig. S1. DRP2A-GFP dynamics in *drp2a-/-drp2b-/-* background.** (A) TIRF-M image of a cell surface of root epidermal cell expressing DRP2A-GFP in *drp2a-/-* background and (B) a representative kymograph of DRP2A lifetime on the PM. Scale bar: 5 µm. (C) TIRF-M image of a cell surface of root epidermal cell expressing DRP2A-GFP in *drp2a-/-;drp2b-/-* background and (D) a representative kymograph of DRP2A lifetime on the PM. Scale bar: 5 µm. (E-G) Data from eleven independent experiments for DRP2A-GFP in *drp2a-/-* and ten independent experiments for DRP2A-GFP in *drp2a-/-;drp2b-/-* were combined to generate a (J) mean recruitment profile of DRP2A-GFP in *drp2a-/-* and DRP2A-GFP in *drp2a-/-;drp2b-/-* foci, (K) mean lifetime (DRP2A-GFP *drp2a-/-*, 26.65±1.13 s; DRP2A-GFP *drp2a-/-;drp2b-/-*, 21.55±0.2 s), and (L) mean density (DRP2A-GFP *drp2a-/-*, 65.23±2.1 spots ROI$^{-1}$; DRP2A-GFP *drp2a-/-;drp2b-/-*, 75.99±4.6 spots

ROI$^{-1}$) of CME events. Plots indicate Mean±SEM DRP2A-GFP *drp2a-/-*, n=11 cells from independent roots, 130,535 tracks; DRP2A-GFP *drp2a-/-;drp2b-/-*, n=10 cells from independent roots, 88,646 tracks. Plot, Mean ± SEM, ***P < 0.0002; *P < 0.0432, t-test to compare to control.

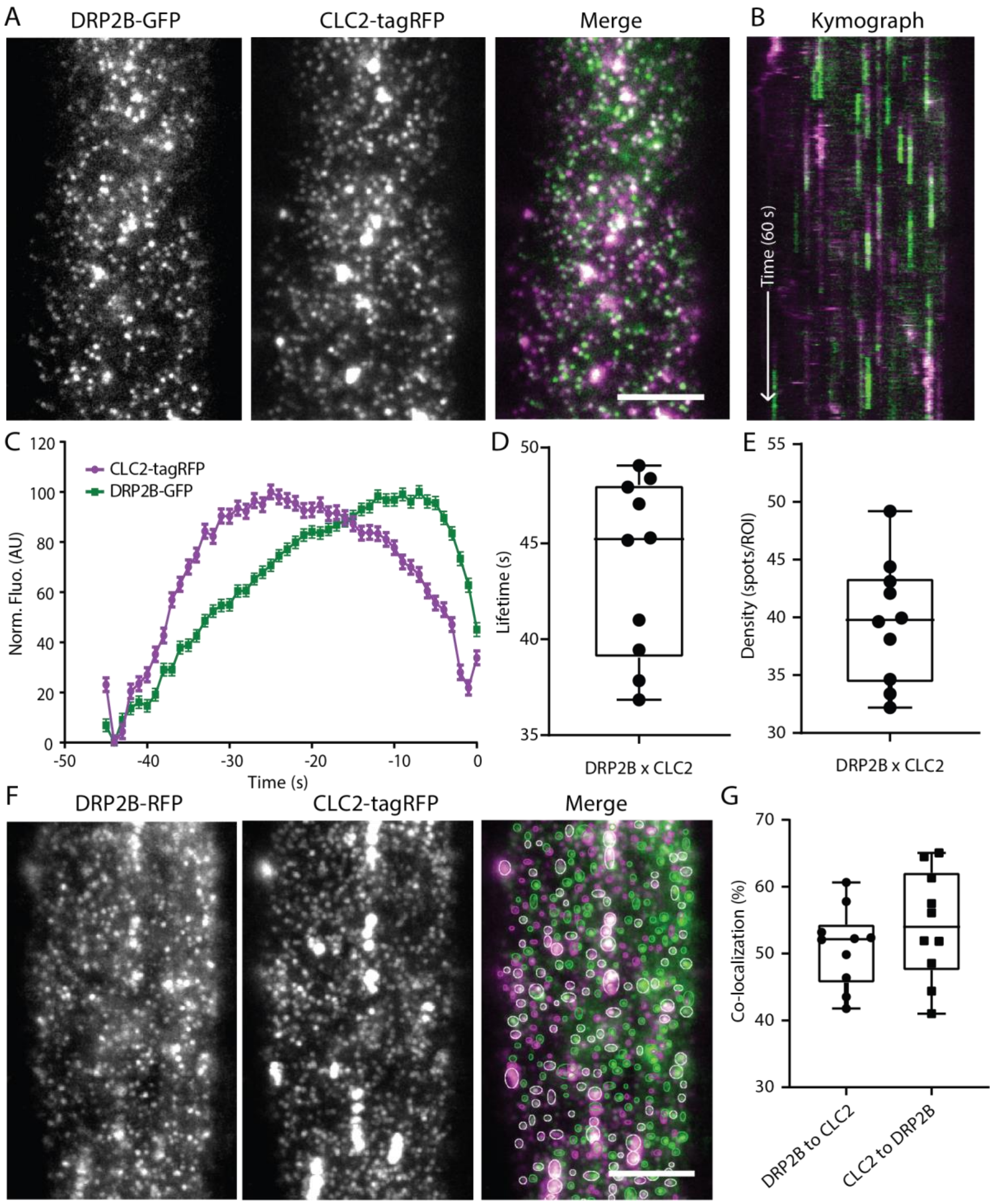


**Fig. S2. Dynamics of DRP2B on the PM.** (A) TIRF-M images of a cell surface of root epidermal cell expressing fluorescently tagged DRP2B-GFP and CLC2-tagRFP. Scale bar: 5 µm. (B) Representative kymograph of DRP2B and CLC2 lifetimes on the PM. (C-E) Data from ten independent experiments were combined to generate a (C) mean recruitment profile of DRP2B to the site of endocytosis, (D) mean lifetime 43.01±0.18 s, and (E) mean density 39.66±1.6 spots ROI$^{-1}$of CME events. Plots indicate Mean±SEM, n=10 cells from independent roots, 27,640 tracks.

(F) Representative image of co-localisation analysis of DRP2B and CLC2 foci. Scale bar: 5 µm.

(G) Quantification of co-localised spots. 50.99±1.8% of DRP2B were co-localised to CLC2, and 54.22±2.5% of CLC2 were co-localised to DRP2B.

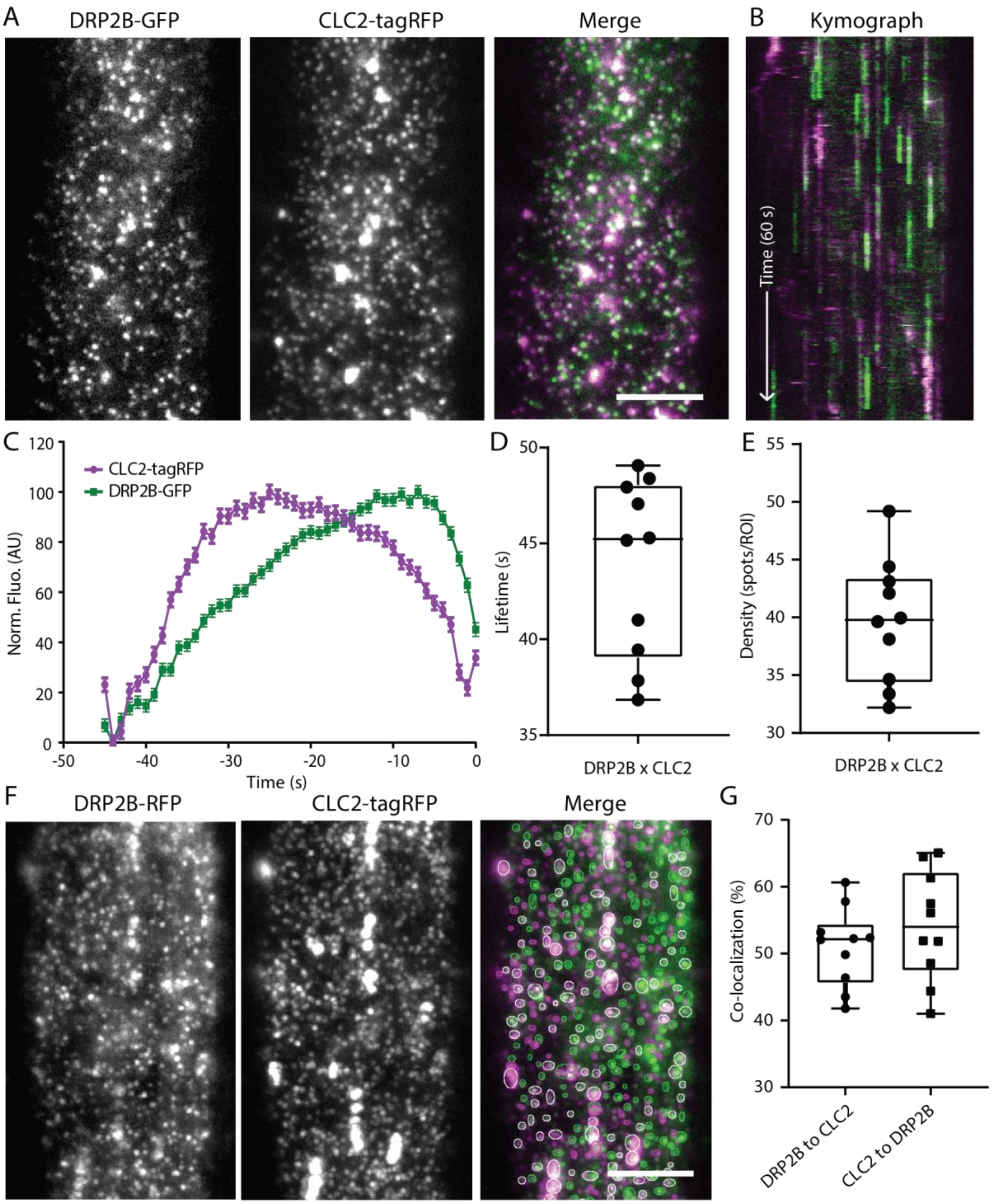


**Fig. S3. SH3P2 characterization *in vitro*.** (A) SDS-PAGE gel of the purified full-length SH3P2 protein. (B) Mass photometry (MS) analysis of FL SH3P2 protein in final concentrations of 50 nM, 75 nM, and 100 nM. Histogram shows the population distributions of purified FL SH3P2. Estimated molecular weights for the monomer and dimer and even counts are indicated on each graph. (C) Representative images of co-sedimentation assay testing lipid-binding capacity of FL SH3P2 to LUVs with different lipid composition: DOPC (100, mol%), DOPC:DOPS (80:20, mol%), DOPC:DOPS:PA (80:18:2, mol%), DOPC:DOPS: $PI(4,5)P_2$ (80:17.5:2,5 mol%). S-supernatant; P-pellet. (D) Quantification of normalised band intensity. Plot, Mean ± SEM, ***$P < 0.0002$; *$P < 0.0269$, t-test to compare to control.

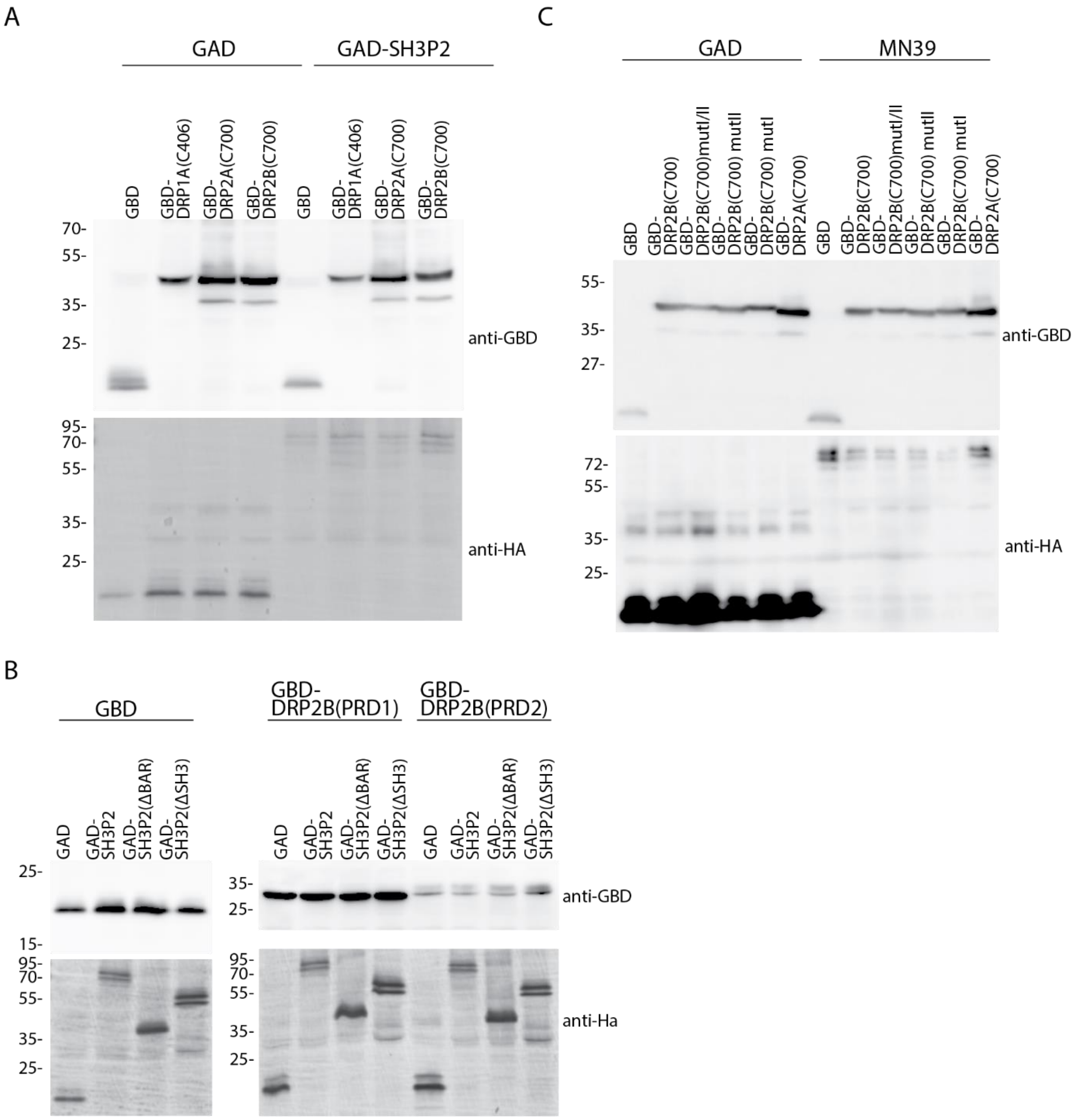


**Fig. S4. Expression of YTH constructs.** (A, B, C) Total proteins were extracted from yeast cells used for YTH analyses in Fig. 4B and subjected to immuno- blotting using anti-GAL4BD and an anti-HA antibody. The anti-HA antibody was used for the detection of GAD-fusion proteins that also contain an HA-tag.

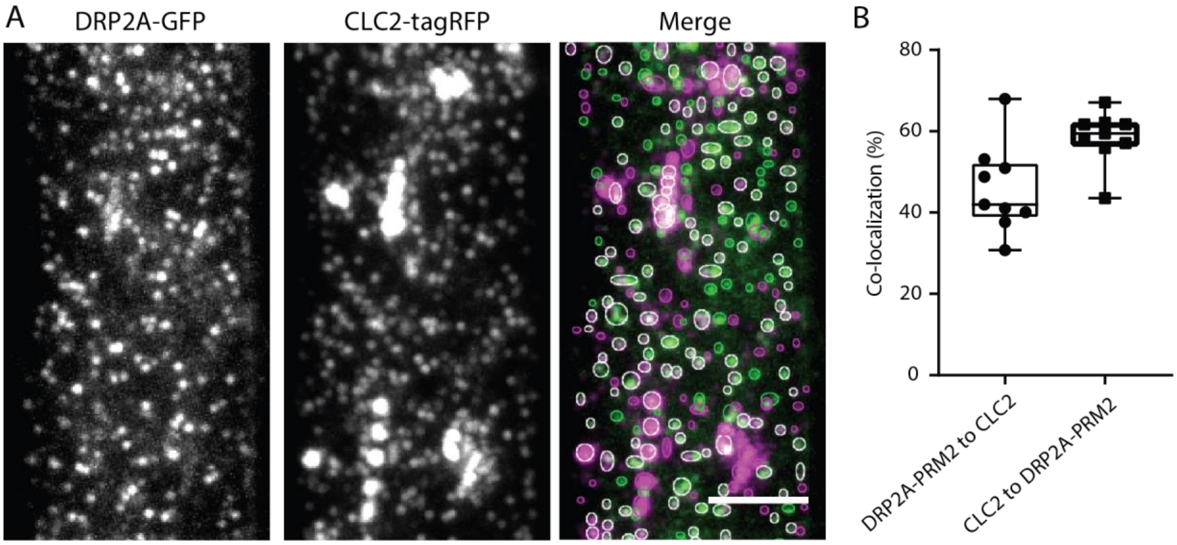


**Fig. S5. Co-localization of DRP2A-PRM2 x CLC2.** (A) Representative TIRF-M image of co-localisation analysis of DRP2A-GFP and CLC2-tagRFP foci. Scale bar: 5 µm. (B) Quantification of co-localised spots. 45.81±3.6% of DRP2A were co-localised to CLC2, and 58.5±2.17% of CLC2 were co-localised to DRP2A.

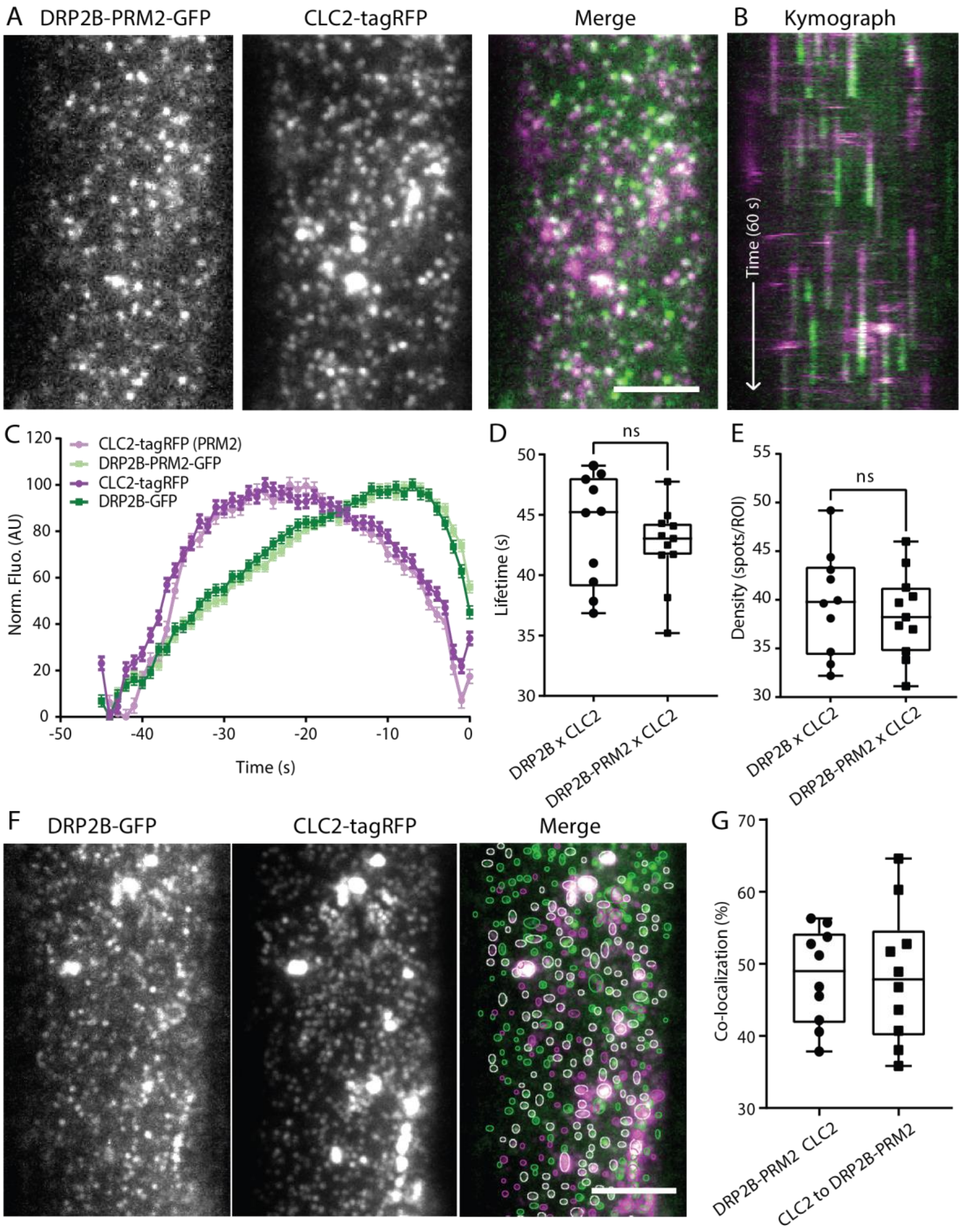


**Fig. S6. Dynamics of DRP2B-PRM2 x CLC2 *in vivo*.** (A) TIRF-M images of a cell surface of root epidermal cell expressing fluorescently tagged DRP2B-PRM2-EGFP and CLC2-tagRFP. Scale bar: 5 µm. (B) Representative kymograph of DRP2B-PRM2 and CLC2 lifetimes on the PM. (C-E) Data from eleven independent experiments for DRP2B-PRM2 x CLC2 were combined and ten independent experiments for DRP2B x CLC2 to generate a (C) mean recruitment profile of

DRP2B and DRP2B-PRM2 foci, (D) mean lifetime (DRP2B, 43.81±1.45 s; DRP2B-PRM2, 42.41±1.01s), and (E) mean density lifetime (DRP2B, 39.67±1.67 spots $ROI^{-1}$; DRP2B-PRM2, 38.49±1.31 spots $ROI^{-1}$) of CME events. Plots indicate Mean±SEM, DRP2B x CLC2, n=10 cells from independent roots, 27,647 tracks; DRP2B-PRM2 x CLC2, n=11 cells from independent roots, 36,486 tracks. Plot, Mean ± SEM, ns >0.05, t-test to compare to control. (F) Representative TIRF-M image of co-localisation analysis of DRP2B-PRM2-GFP and CLC2-tagRFP foci. Scale bar: 5 µm. (G) Quantification of co-localised spots. 48.26±2.1% of DRP2B-PRM2 were co-localised to CLC2, and 48.32.5±2.9% of CLC2 were co-localised to DRP2B-PRM2.

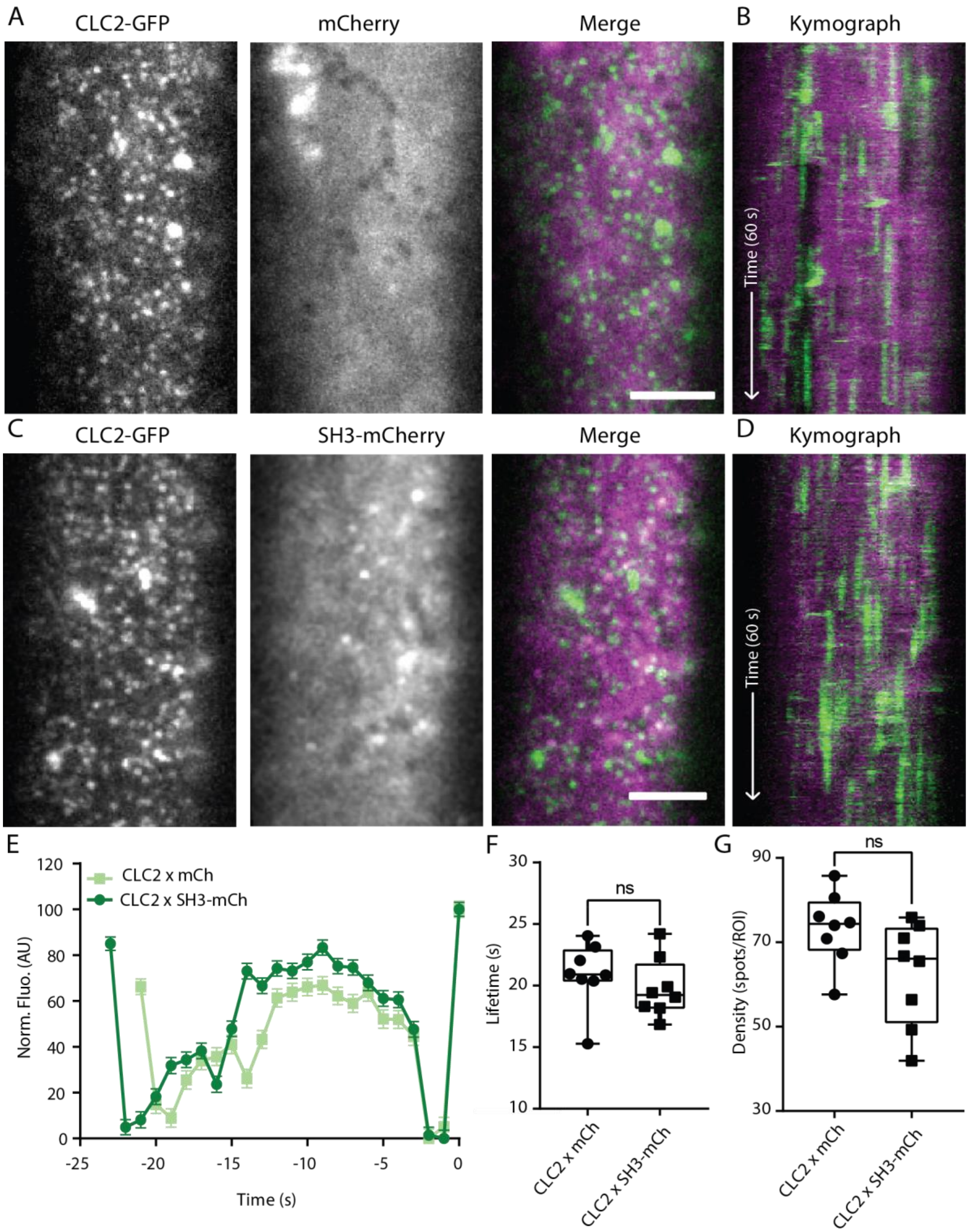


**Fig. S7. Overexpression of the SH3 domain does not influence the dynamics of CLC2 *in vivo*.** (A) TIRF-M images of a cell surface of root epidermal cell expressing fluorescently tagged CLC2-EGFP and free mCherry. Scale bar: 5 µm. (B) Representative kymograph of CLC2 lifetimes on the PM after 24h mCherry overexpression. (C) TIRF-M images of a cell surface of root epidermal cell expressing fluorescently tagged CLC2-EGFP and SH3-mCherry of SH3P2. Scale bar: 5 µm. (D) Representative kymograph of CLC2 lifetimes on the PM after 24h SH3-mCherry overexpression. **(**E-G) Data from eight independent experiments for CLC2 x mCherry and eight independent experiments for CLC2 x SH3-mCherry were combined to generate a (C) mean recruitment profiles of CLC2 foci, (D) mean lifetime (CLC2 x mCherry, 20.9±0.92 s; CLC2 x SH3-mCherry, 19.78±0.8s), and (E) mean density lifetime (CLC2 x mCherry, 73.38.67±3 spots $ROI^{-1}$; CLC2 x SH3-mCherry, 62.58±4.3 spots $ROI^{-1}$) of CME events. Plots indicate Mean±SEM, CLC2 x mCherry, n=8 cells from independent roots, 91,321 tracks; CLC2 x SH3-mCherry, n=8 cells from independent roots, 40,268 tracks. Plot, Mean ± SEM, ns >0.05, t-test to compare to control.

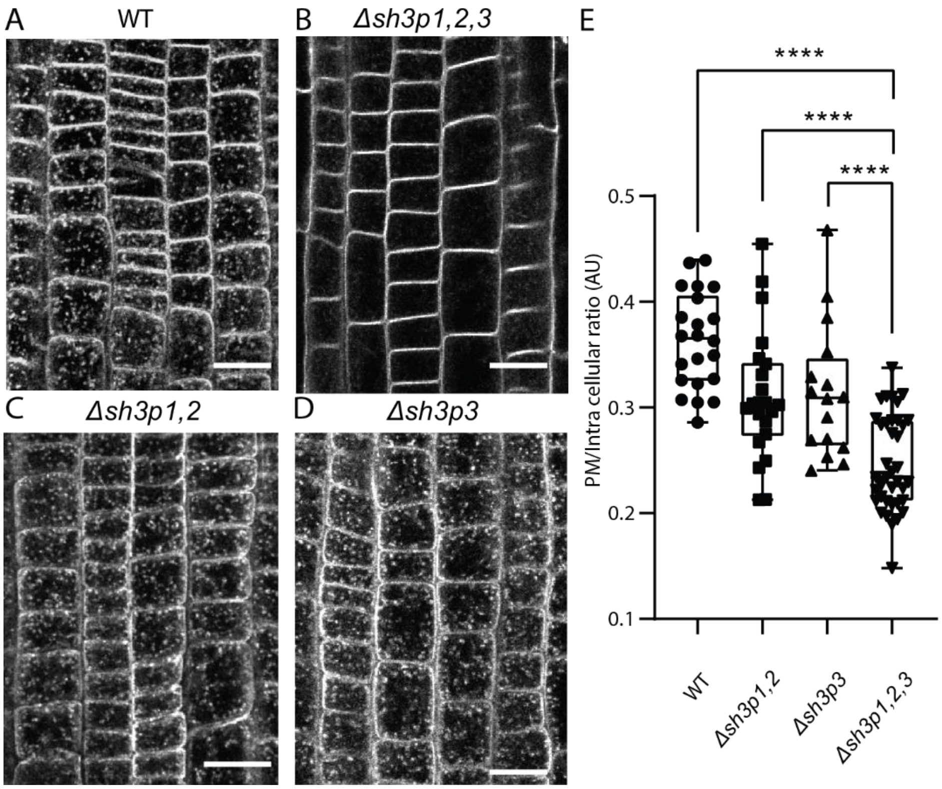


**Fig. S8.** ***Δsh3p1,2,3*** **triple mutant has reduced membrane uptake.** (A-D) Confocal images of epidermal root cells of *A.thaliana* Col-0 (WT), *Δsh3p1,2, Δsh3p3, Δsh3p1,2,3,* treated with 2 µM membrane dye FM4-64 15 min, RT. Scale bar: 20 µm. (E) Quantification of the membrane uptake. Plots indicate Mean±SEM. For WT, n=22 independent seedlings, 697 cells; for *Δsh3p1,2*, n=22 independent seedlings, 582 cells; *Δsh3p3,* n=16 independent seedlings, 417 cells; *Δsh3p1,2,3,* n=35 independent seedlings, 1060 cells. Plot, Mean ± SEM, ****P < 0.001, t-test to compare to control.

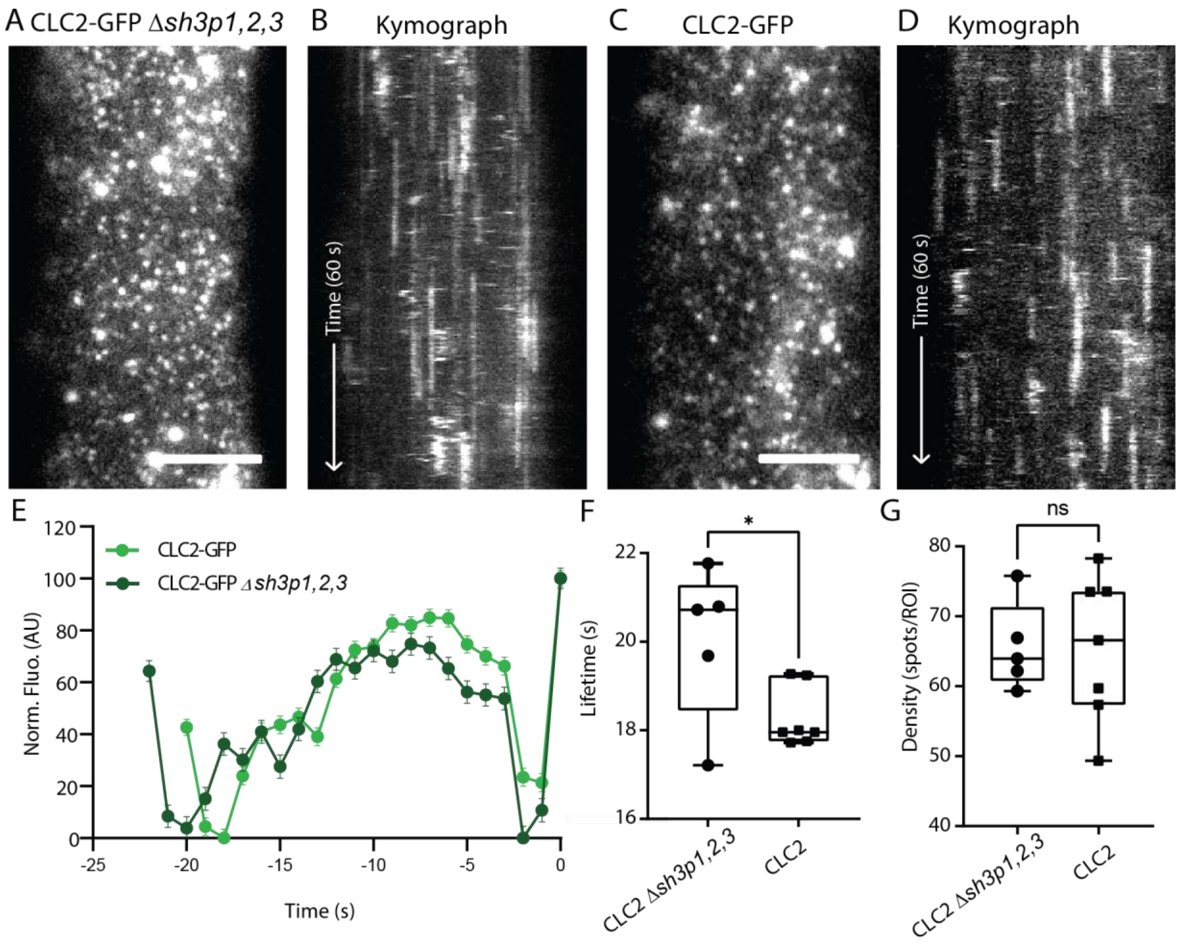


**Fig. S9. Dynamics of CLC2-GFP in *Δsh3p1,2,3* mutant.** (A) TIRF-M image of a cell surface of root epidermal cell expressing CLC2-GFP in *Δsh3p1,2,3* background and (B) a representative kymograph of CLC2 lifetime on the PM. (C) TIRF-M image of a cell surface of root epidermal cell expressing CLC2-GFP and (D) a representative kymograph of CLC2-GFP lifetime on the PM. Scale bar: 5 µm. (E-G) Data from seven independent experiments for CLC2-GFP and five independent experiments for CLC2 *Δsh3p1,2,3* were combined to generate a (E) mean recruitment profile of CLC2 and CLC2 *Δsh3p1,2,3* foci, (F) mean lifetime (CLC2, 18.27±0.25 s; CLC2 *Δsh3p1,2,3,* 20.04±0.77 s), and (G) mean density (CLC2, 65.48±3.9 spots ROI$^{-1}$; CLC2 *Δsh3p1,2,3,* 65.63±2.8 spots ROI$^{-1}$) of CME events. Plots indicate Mean±SEM CLC2, n=7 cells from independent roots, 46,995 tracks; CLC2 *Δsh3p1,2,3,* n=5 cells from independent roots, 44,205 tracks. Plot, Mean ± SEM, *P < 0.0432, ns>0.05, t-test to compare to control.

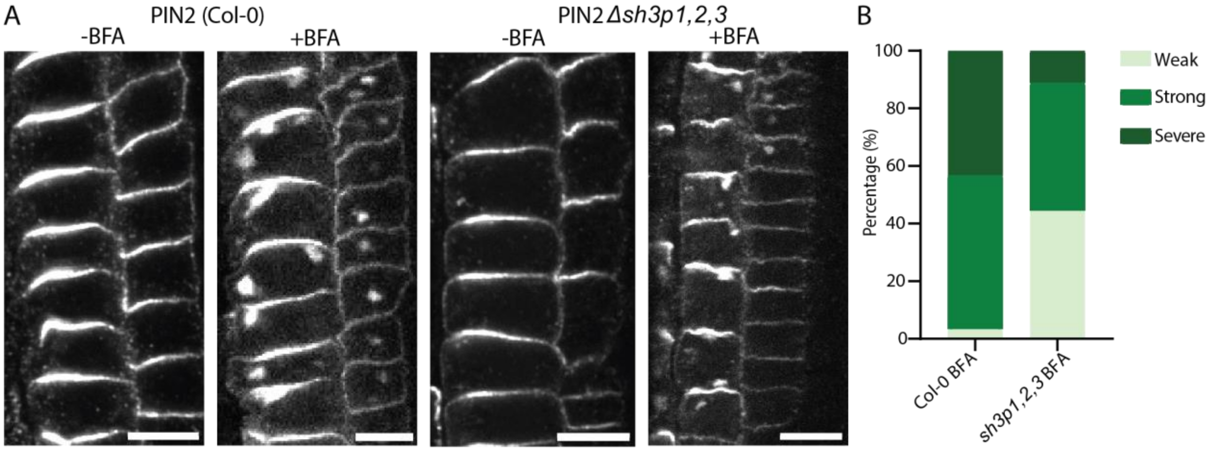


**Fig. S10. General dynamics of PIN2 proteins in *Δsh3p1,2,3* triple mutant.** (A) Confocal images of PIN2 localization in Epidermis E and Cortex C cells of *Arabidopsis* roots untreated or treated with 50 µM BFA for 1 h in Col-0 or *Δsh3p1,2,3* triple mutant. Scale bar 20 µm. (B) A stacked column chart representing the percentage of roots, per genotype, distributed in three different categories (weak, strong, severe) based on the abundance of PIN2 BFA bodies (internalization) formed in the cells after BFA treatment.
